# “A CRISPR-dCas13 RNA-editing tool to study alternative splicing”

**DOI:** 10.1101/2022.05.24.493209

**Authors:** Yaiza Núñez-Álvarez, Tristan Espie--Caullet, Géraldine Buhagiar, Ane Rubio-Zulaika, Josune Alonso-Marañón, Elvira Perez-Luna, Lorea Blazquez, Reini F. Luco

## Abstract

Alternative splicing allows multiple transcripts to be generated from the same gene to diversify the protein repertoire and gain new functions despite a limited coding genome. It can impact a wide spectrum of biological processes, including disease. However, its significance has long been underestimated due to limitations in dissecting the precise role of each splicing isoform in a physiological context. Furthermore, identifying key regulatory elements to correct deleterious splicing isoforms has proven equally challenging, increasing the difficulty to tackle the role of alternative splicing in cell biology. In this work, we take advantage of dCasRx, a catalytically inactive RNA targeting CRISPR-dCas13 ortholog, to efficiently switch alternative splicing patterns of endogenous transcripts without affecting overall gene expression levels in a cost-effective manner. Additionally, we demonstrate a new application for the dCasRx splice-editing system to identify key regulatory RNA elements of specific splicing events. With this approach, we are expanding the RNA toolkit to better understand the regulatory mechanisms underlying alternative splicing and its physiological impact in various biological processes, including pathological conditions.

## INTRODUCTION

More than 90% of human multi-exon genes are alternatively spliced into different mature mRNAs to increase transcript diversity (1). In this process, exonic and intronic sequences can be included or excluded via exon skipping, intron retention, the use of mutually exclusive exons and alternative 3’ or 5’ splice sites (2). Such RNA rearrangements can introduce premature STOP codons that lead to protein truncation or induction of nonsense-mediated mRNA decay (3). They can also modify protein domains important for cellular localization, protein-protein interaction, regulation of enzymatic activity and DNA/RNA binding amongst others, increasing like this the cell’s proteome diversity (4, 5). Unfortunately, such transcript plasticity can also lead to disease, with around 15% of genetic diseases due to mutations in splicing-related genes. Importantly, splicing can now be modulated with innovative therapeutic approaches, as for the already approved spinal muscular atrophy (SMA) or Duchenne muscular dystrophy (DMD) (6–8).

While improvements in RNA sequencing technologies and mass spectrometry sensitivity have exponentially increased the number of identified splicing isoforms, their function and biological impact remain rather unknown with the misconception that small changes in alternative splicing at the molecular level might be irrelevant at the cellular level (5). Several approaches have aimed at exogenously expressing, or selectively downregulating, the splicing isoform of interest to address its functional role using expression vectors, or splice junction-specific short hairpin shRNA and small interfering siRNA, respectively. However such strategies have often ended up with misexpression of the targeted gene at non-physiological levels, with the subsequent overexpression or knock-down of the studied protein, which does not reflect a change in splicing, but a change in the overall gene expression levels of the studied gene (9, 10). Splice-switching antisense oligonucleotides (SSO) can induce a change in the splicing outcome, shifting splicing isoforms without impacting overall expression levels (11). However, this is an expensive approach that requires the screening of several oligonucleotides at high quantities for an efficient and time-sustained splicing effect, which reduces its feasibility in large-scale studies (11). Small molecule splicing modulators, which are synthetic analogues to inhibit splicing reactions, are an innovative new strategy of special interest in the clinic (12). Unfortunately, there are difficult to implement and can lead to many indirect effects.

Recently, with the discovery of a unique prokaryotic immune system to eliminate nucleic acid fragments from bacteriophages, called the Clustered Regularly Interspaced Short Palindromic Repeats (CRISPR) (13), key cis-regulatory splicing regions can now be targeted at the DNA level to cause a point mutation (or deletion) at a specific regulatory region that will impact splicing (14–16). However, this approach is limited to the existence of a protospacer adjacent motif (PAM) near the targeted region for Cas9 cleavage, and it requires the modification of both alleles in an irreversible way.

The discovery in 2016 of a new family of RNA-targeting CRISPR-associated nucleases, called Cas13, was a game-changer for RNA editing (17, 18). Cas13, and their catalytically inactive dCas13 mutants, are RNA-guided RNases that comprise at least 6 subfamilies: (d)Cas13a, b, c, d, X and Y (19). RfxCas13d (from *Ruminococcus flavefaciens*), also named CasRx, is amongst the most efficient Cas13 RNAses, together with PspCas13b (from *Prevotella sp*.) and PguCas13b (from *Porphyromonas gulae)* (20–26). These protein effectors have proven useful for not only downregulating specific mRNAs *in cellulo* (*17, 21, 22, 25, 26*) and *in vivo* in different model organisms (24, 27, 28), but also for viral RNA detection (28, 29), site-directed RNA editing (19, 20, 30), modulation of m^6^A levels (31, 32), RNA live imaging (33) and interference of alternative polyadenylation sites (23). In contrast to Cas9, Cas13b and Cas13d family members have no targeting sequence constraints, such as a protospacer flanking site (PFS) or protospacer adjacent motif (PAM), allowing any sequence in the transcriptome to be potentially targeted (20, 21). Furthermore, despite some degree of target-dependent collateral ribonuclease activity, mostly when targeting exogenous overexpressed genes in contrast to transiently targeted endogenous genes (17, 19, 25), these RNAses have greatly reduced off-target effects compared to other gene-specific targeting systems (such as RNA interference), and with no impact on the genome (20, 21), increasing its interest as a versatile editing tool.

Regarding RNA splicing, dCasRx is the most widely used dead dCas13 to modulate alternative splicing. However, most of the studies are focused on exogenously expressed splicing reporters, or have used dCasRx proteins fused to a splicing regulator, which could increase indirect effects (4, 21, 34). In this work, we are addressing important aspects of unconjugated dCas13-mediated splicing editing to establish a user-friendly guideline of how to induce physiologically relevant changes in the alternative splicing of endogenous genes. Furthermore, limitations for RNA splicing editing are discussed, and a new application for dCasRx is proposed in the identification of key RNA regulatory regions for functional editing and study of the splicing regulators involved.

## MATERIAL AND METHODS

### Cell culture of human cell lines

Human Embryonic Kidney (HEK) 293T and HeLa cells were maintained at 37°C with 5% CO2 in DMEM (4.5 g/L glucose) with Glutamax, supplemented with 10% FBS (Heat inactivated Sigma F9665-500ML) and 1% penicillin/streptomycin. Cells were maintained in subconfluent conditions and passaged at 1:10 ratio every 3 days. HEK293T cell line was a gift from Dr. Nadine Laguette at IGMM and HeLa from Dr. Mounira Amor-Gueret at Institut Curie. The cells have been authenticated and periodically tested for mycoplasma.

### Cloning and Plasmids

To allow indirect quantification of gRNA transfection via mCherry expression, pXR003 processed gRNA (Addgene #109053) was subcloned in pKLV2.4-Hygro mCherry gRNA lentiviral plasmid (35) using EcoRI and Mlu restriction enzymes (FW: CCCACGCGTGAGGGCCTATTTCCCATGATTC, REV: CCCCCGAATTCAAAAAAAAGGTCTTCTCGAAGACC). The bleomycin resistance cassette of the dCasRx plasmid (pXR002, Addgene #109050) was replaced by a blasticidin resistance cassette. For dCasRx-hnRNPA1 fusion protein, the Gly-rich C-terminal effector domain of hnRNPA1 (from 196 to 320 amino acids) was added to dCasRx as in (21). Briefly, hnRNPA1 effector domain was amplified from HEK293T cDNA using the Q5 High fidelity polymerase (#M0491S, NEB). Overhanging BsWI sites were added to the oligonucleotides for cloning into the dCasRx plasmid with Quick Ligation kit (M2200S, New England Biolabs) (FW: gtgacgtacggccaccatgGGTCGAAGTGGTTCTGG, REV: catggtggccgtacgtccAAATCTTCTGCCACT).

### Transient transfection of human cell lines

For splicing and RNA abundance analysis, HEK293T and HeLa cells were plated at a density of 10,000 cells per well in a 96-well plate. Transfection with 200 ng of (d)CasRx or other (d)Cas13 expression plasmids and 200 ng of the indicated gRNA was done with Lipofectamine 2000 (Life Technologies) according to the manufacturer’s protocol. HEK293T cells were harvested 72 hours post-transfection. HeLa cells were selected for dCasRx expression with 20 ng/μL blasticidin for a total of 48 hours post-transfection

For competition experiments, HEK293T cells were plated at a density of 10,000 cells per well in a 96-well plate and transfected with 200 ng of dCasRx expression vector, 200 ng of *CTNND1.*ex2 (gRNA_e2) or *PKM*.ex10 (CD) targeting gRNA and 200 ng of *SRSF1* or *SRSF3* expression vector using Lipofectamine 2000 (Life Technologies). Non-targeting (NT) gRNA and empty expression vector were used as controls. Transfected cells were harvested 72 hours post-transfection.

For protein expression analysis, HEK293T cells were plated at a density of 200,000 cells per well in a 24-well plate and transfected with 800 ng of dCasRx expression plasmid and 800 ng of gRNA using Lipofectamine 2000 (Life Technologies). Transfected cells were harvested 72 hours post-transfection.

### gRNA design

For custom tiling gRNA design, no special requirements were considered but being complementary to the RNA transcript to be edited. When using the “cas13design” software (https://cas13design.nygenome.org, gRNAs are annotated as CD), the most abundant transcript isoform expressed in HEK293T, including the alternatively spliced exon of interest, was selected for gRNA design. Top-scored gRNAs were used for splicing editing. When using the CHOPCHOP software (https://chopchop.cbu.uib.no/, gRNAs are annotated as CC), we used the *Homo sapiens* hg38 genome version and the option “CRISPR/Cpf1 or CasX” without any PAM requirements.

In all cases, gRNAs were annotated from acceptor to donor. When both intronic and exonic regions were targeted with different gRNAs along the same pre-mRNA, the gRNAs were labelled as “iX” for intronic and “eX” for exonic. gRNAs located in acceptor and donor regions were labelled as “Acc” or “Don”, respectively.

### gRNA cloning

gRNAs were ordered as desalted oligonucleotides at 100 µM at Eurofins Genomics with the corresponding overhanging BbsI complementary sites as indicated in Supplementary Table 2 for (d)CasRx gRNAs and in Supplementary Table 3 for (d)Cas13b gRNAs. 100 nmol of forward and reverse gRNA oligonucleotides were annealed in NEB Buffer 4 (Vf=50μL) by heating at 98 °C for 5 min and progressive cooling down to room temperature in the thermoblock for ∼3h. 3µl of annealed oligonucleotides were cloned in 50ng of gRNA backbone vector (previously cut with BbsI-HF and gel extracted) using Quick Ligation kit (#M2200S, New England Biolabs) and transformed in NEB 5-alpha Competent E. coli (C2987I, New England Biolabs). Positive colonies were screened by PCR using GoTaq G2 Hot Start green Master Mix (#M7422, Promega) using the consensus pLKO F oligonucleotide (5’->3’GACTATCATATGCTTACCGT) and the corresponding reverse primer used to clone the gRNA, in a final volume of 12,5 µl using the following program: 95°C_2min, 30x(95°C_30”, 54°C_30”, 72°C_10”), 72°C_2min.

### shRNA cloning and splicing factor knock-down

shRNAs were ordered as oligonucleotides with the corresponding overhanging sequences (Supplementary Table 5), for annealing and cloning in digested pLKO.1-blast (with AgeI-HF and EcoRI-HF) as explained in “gRNA cloning”.

For SRSF1, MBNL1, ELAV1, CELF1, KHDRBS3, PTBP1 and FUS knock-down, HEK293T cells were plated at 2.5×10^6^ cells in a 100mm dish. Cells were transfected with 5µg of the shRNA plasmid and 250mM Cacl2 in 500µL sterile water. Samples were gently mixed and completed with 2X HEPES Buffered Saline (HBS) for 10 min incubation at room temperature. The shRNA mix was gently dropped on HEK293T cells and medium was replaced 16h after transfection. RNA was extracted 72h post-transfection with ThermoFisher’s GenJet RNA purification kit (#K0732). For SRSF3 and hnRNPA1 knock-down, HEK293T cells were plated in 96 well plates and transfected with 200ng of shRNA plasmid as explained in “Transient transfection of human cell lines”. Cells were collected 72h after transfection and RNA was extracted using TurboCapture 96 mRNA Kit (#72251, QIAGEN) following manufacturer’s protocol. Mission SHC002 non-mammalian shRNA was used as control (Sigma-Aldrich).

### Gene expression and alternative splicing RT-qPCR analysis

Cells were lysed by directly adding 35 μl of β-mercaptoethanol-supplemented TCL buffer for mRNA extraction using TurboCapture 96 mRNA Kit (#72251, QIAGEN) following manufacturer’s protocol. Captured mRNA was eluted in 11 μl of TCE buffer and incubated at 65°C for 5 min. Eluted mRNA was then reverse transcribed using random hexamer primers and Transcriptor First Strand cDNA Synthesis Kit (#04897030001, Roche) at 65°C for 10 min, 25°C for 10 min, 50°C for 60 min, and 85°C for 5 min. cDNA was diluted 1/5 and quantified by RT-qPCR in 10 μL final reactions using 2X iTaq Universal Sybr Green Supermix (#1725125, Bio-Rad) in CFX96 Touch Real-Time PCR machines (Bio-Rad). The oligonucleotides used are listed in Supplementary Table 4. To avoid DNA amplification when quantifying changes in RNA splicing, oligos were designed with IDT Realtime PCR Tool and Primer3 software spanning an intronic region. When studying changes in total gene expression levels, we amplified constitutive exons far from the targeted RNA region to properly address knock-down efficiency of the whole mRNA molecule and not just the targeted region. Gene expression levels were normalized with the universal housekeeping gene *TBP* using the 2^(ΔCt) method. Regarding splicing, the percentage of exon inclusion was calculated using the same method, but relative to total expression levels of the corresponding gene. For evaluation of dCasRx splice-editing efficiency, changes in exon inclusion levels were represented as the fold change respect the levels obtained with transfection of a non-targeting gRNA performed in parallel in the same plate.

### RNA Immunoprecipitation

Transfected HEK293T cells were collected in Lysis buffer (150 mM KCl, 25 mM Tris-HCl pH7.4, 5mM EDTA, 0.5% NP40, 0,5 mM DTT, 100U/mL RNase inhibitor (Promega) and EDTA-free proteinase inhibitor (Roche)). Upon incubation on ice for 15 min, samples were centrifuged at 15,000xg for 15 min at 4°C and the supernatant was pre-cleared with protein G-magnetic Dynabeads (#10004D, Invitrogen) at 4°C for 2h. Meanwhile, and for each condition, 5 µg of anti-SRSF3 (#334200, Invitrogen) or anti-IgG (#Sc-2025, Santa Cruz) antibody was pre-incubated with protein G-Dynabeads in Lysis buffer at 4°C for 2h. Control IgG and SRSF3 antibody-coated beads were then incubated with pre-cleared lysates overnight at 4°C in a rotating wheel. 10% input was taken before incubation with the beads. Beads were then washed three times with lysis buffer. Proteins were extracted with SDS-PAGE loading buffer (#NP0007, FisherSci) at 70°C during 10 min while mixing at 1100 rpm for western blot validation of SRSF3 immunoprecipitation. RNA was purified with TRIzol Reagent (#10296010, Fisher Scientific) using phase lock gel tubes (#733-2478, VWR) following manufacturer’s instructions. Briefly, upon 5 min incubation at room temperature while mixing at 1,100 rpm, chloroform was added, samples were centrifuged at 12,000xg for 20 min at 4°C and the top layer was collected for RNA precipitation overnight at-20°C with isopropanol, 200mM sodium acetate and glycoblue (#AM9516, Fisher Scientific). RNAs were then treated with DNAse I (#EN0521, ThermoFisher) and reverse-transcribed with High-Capacity cDNA kit (#4368814, Fisher Scientific). The primers used in real time qPCRs are listed in Supplementary Table 4. All RNA-IP Ct values were compared to Input Ct and further normalized to either the values obtained with IgG to evaluate SRSF3 IP efficacity, or to the values obtained at SRSF3 transcript, used as a positive control, to reduce intra-experimental variability.

### Agarose gel PCR and Long-read amplicon sequencing for splicing analysis

Total RNA from dCasRx+gRNA transfected HEK293T cells was reverse transcribed using oligo dT and the Transcriptor First Strand cDNA Synthesis Kit (#04897030001, Roche) as manufacturer’s recommendations. *CD46* or *PKM* whole transcript cDNA was amplified using the Q5 High fidelity polymerase (#M0491S, NEB) and the oligonucleotides indicated in Supplementary Table 4 using the program: 98°C_30’’, 40x(98°C_10”, 68°C_30”, 72°C_2min), 72°C_5min. Some of the *CD46* PCR product was run on 2% agarose gel for separation of long PCR products with small differences in size (93b), while *PKM* PCR product was run on a 0.8% agarose gel. The rest of the PCRs were purified with the NucleoSpin Gel and PCR Clean-up XS kit (#740611.50, Macherey-Nagel) for long-read amplicon sequencing using Oxford Nanopore Technology (via Plasmidsaurus).

ONT Nanopore full-length transcript FASTQs reads were then aligned to the human reference genome (GRCh38) using minimap2 (version 2.26-r1175, https://doi.org/10.1093/bioinformatics/bty191) in the splice-aware mode (-ax splice) and providing a junction reference BED file (gencode v41,--junc-bed). BAM files were then sorted and indexed with samtools (version 1.17). In order to visualize the splicing events, ggsashimi was used filtering for events with more than 6 reads (version 1.1.5, https://github.com/guigolab/ggsashimi).

### Western blot of PKM isoforms

Transfected cells from 24-well plates were lysed in RIPA buffer containing 1X protease cOmplete inhibitors (#11836145001, Sigma) and quantified using BCA method (BCA Protein Assay, 23227, Pierce). 40 μg of protein/sample were run at 200mV for 1h30 using XCell SureLock Mini-Cell System (ThermoFisher). NuPAGE 4-12% Bis Tris gels (#NP0322B0X, Invitrogen) were transferred in the same system at 65mV for 1h. Membranes were cut according to protein weight to allow the analysis of the target isoforms and loading controls in the same membrane. Membranes were then blocked with 5% w/v BSA in 1x TBS-Tween for 1h and then incubated with the corresponding primary antibodies diluted in blocking buffer o/n at 4°C with shacking. After incubation with the appropriate secondary antibodies, protein levels were semi-quantified with the ECL Supersignal West Pico Chemiluminescent Substrate (#34080, ThermoFisher) and the Chemidoc Gel Imaging System (Bio-rad) following manufacturer’s recommendations.

### RNA Motif search analysis

RNA binding motif search analysis was performed using *CTNND1* exon2 sequence in four public software: RBPDB v1.3 (http://rbpdb.ccbr.utoronto.ca), RBPMAP v1.1 (http://rbpmap.technion.ac.il), SFMAP v1.8 (http://sfmap.technion.ac.il/) and Spliceaid (http://www.introni.it/splicing.html). Software was run with default parameter settings, except for RBPDB, in which a threshold of 0.8 was applied. For RBPMAP, a HIGH Stringency level was applied at all the motifs available in Human and Mouse. For SFMAP both “Perfect match” and “High stringency” levels were selected. Only the motifs predicted by at least two of the four software were kept.

### Statistical Analysis

All statistical analyses were performed using GraphPad Prism 9. Data were represented as means ± SD of at least three independent experiments, in which each individual replicate was plotted as a dot. Statistical analysis was carried out using Student’s *t* test for comparing two conditions, or one-way ANOVA for multiple comparisons, as in the tiling of several gRNAs across an exon. A *P* value less than 0.05 was considered to be statistically significant. (\**P* < 0.05; \*\**P* < 0.01; \*\*\**P* < 0.001; \*\*\**P* < 0.0001; ns, no significance).

## RESULTS AND DISCUSSION

### A position-dependent effect of dCasRx-mediated modulation of splicing

It was previously shown that targeting catalytically inactive dCasRx to regulatory splice sites flanking the alternatively spliced exon, such as the branch point or the donor and acceptor splice sites, could induce exon skipping (21). Best splice-editing results were obtained when using a combination of gRNAs targeting several regions along the exon or by fusing to dCasRx the effector domain of a splicing regulator, such as hnRNPA1 (21) or RBFOX1 (34). Since most of these gRNAs were tested in splicing reporters, we aimed at elucidating the impact of unfused dCasRx in the alternative splicing of endogenous genes.

We first targeted dCasRx, or the catalytically active CasRx endonuclease, to an alternative splicing event intimately involved in cancer, which is the exon 2 of the catenin delta-1 protein (CTNND1) (36, 37). Exon 2 inclusion changes the translation-initiation start site of *CTNND1*, which impacts its capacity to interact with E-cadherins, causing destabilization of cell-cell interactions and an increase in cell motility (38). Furthermore, we have previously shown that induction of *CTNND1* exon 2 inclusion, using dCas9 epigenome editing tools, is sufficient to increase the migration and cell invasiveness of normal human epithelial MCF10a cells (35), highlighting the interest of developing efficient tools to reduce exon 2 inclusion in highly invasive cancer cells. To identify the gRNA with the strongest splice-editing effect in HEK293T cells, we designed a tiling array of 8 individual gRNAs covering the whole exon with a 5nt overlap (Figure 1A and Supplementary Figure S1A).

**Figure 1.**
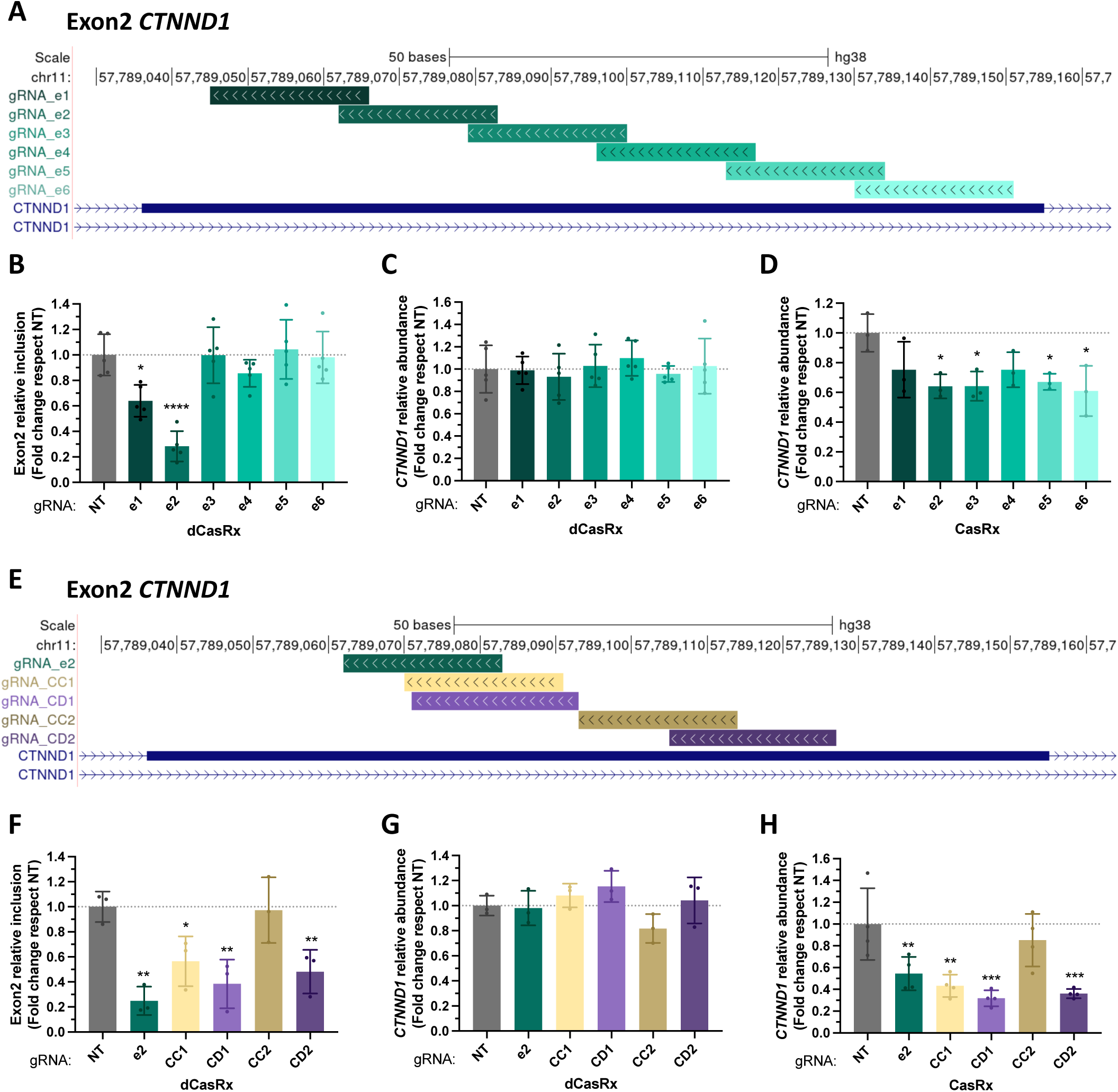
Position-dependent effect of dCasRx-mediated modulation of splicing. (**A**) Genomic location of 6 overlapping gRNAs tiled across *CTNND1* exon 2. (**B**) *CTNND1* exon 2 relative inclusion levels in HEK293T cells after transfection of dCasRx and the indicated gRNAs. (**C,D**) CTNND1 relative RNA abundance after transfection of dCasRx (**C**) or CasRx (**D**) with the indicated gRNAs. (**E**) Genomic location of top-ranked gRNAs targeting *CTNND1* exon 2 using “Cas13design” (CD, purple shades) or CHOPCHOP (CC, yellow shades) web tools. Best performing custom-made splice-editing gRNA_e2 is shown as a control reference (**F**) *CTNND1* exon 2 relative inclusion levels after transfection of dCasRx and the indicated gRNAs. (**G,H**) *CTNND1* relative RNA abundance after transfection of dCasRx (**G**) or CasRx (**H**) and the indicated gRNAs. In all conditions, exon 2 RT-qPCR levels were normalized by *CTNND1* total expression; while *CTNND1* total RNA levels were normalized by *TBP* housekeeping gene expression. Data are represented as mean ± SD of the fold change relative to non-targeting gRNA (NT) in at least 3 biological replicates. *P <0.05, **P <0.01, ***P <0.001, **** P <0.0001 in unpaired one-way ANOVA respect NT.

Contrary to what was published in splicing reporter minigenes, targeting the acceptor or donor splice sites with specific gRNAs had minor impact on *CTNND1* exon 2 splicing (Supplementary Figure S1B). However, one of the gRNAs targeting exon 2 (gRNA_e2) strongly reduced exon inclusion more than 70%, without impacting *CTNND1* total expression levels (Figure 1B,C and Supplementary Figure S1E,F). While the other gRNAs also targeting exon 2 had minor or no impact on the exon’s inclusion levels. To test whether other dCas13 family members could be more efficient editing splicing, we compared dCasRx effect with dPspCas13b or dPguCas13b, which are the two other (d)Cas13 proteins found to efficiently target the RNA for protein knock-down (20, 21, 26), live imaging of RNAs (dPspCas13b fused to EGFP, (33)) or interference with polyadenylation sites (unfused dPguCas13b, (23)). HEK293T cells were transfected with dCas13 orthologs and equivalent gRNA sequences adapted for each dCas13 family member. Despite having an impact on *CTNND1* RNA abundance when the gRNAs were transfected with PspCas13, catalytically-inactive dPspCas13b did not change splicing (Supplementary Figure S1H-J). Neither did dead dPguCas13b ortholog (Supplementary Figure S1K-L), pointing to dCasRx as the dCas13 of choice for splicing editing.

We then evaluated whether the cellular concentration of dCasRx and/or the gRNA could limit splice-editing efficiency, as it needs to target all the pre-mRNA molecules present in the nucleus. To test this, we transfected increasing levels of CasRx, dCasRx and gRNA plasmids in HEK293T cells (Supplementary Figure S1M-U). We used gRNA_e1 and gRNA_e2 as the only gRNAs impacting *CTNND1* splicing. We first confirmed that transfection of the gRNA alone could not impact splicing, nor mRNA levels, supporting a (d)CasRx-mediated effect (Supplementary Figure S1O,R). Increasing CasRx (Supplementary Figure S1M-O), dCasRx (Supplementary Figure S1P-R) or gRNA expression levels (indirectly assessed with mCherry, present in the gRNA plasmid-Supplementary Figure S1S-U) did not significantly improve RNA cleavage nor splicing changes, suggesting that high quantities of CRISPR-associated protein or gRNA are not necessary for efficient RNA editing.

dCasRx can thus efficiently induce isoform-switching splicing changes in an endogenous gene without impacting overall gene expression levels. However, not all the gRNAs tested had the same effect on *CTNND1* exon 2 splicing, raising the question how to design a strong splice-editing gRNA (Figure 1B). Turns to that based on CasRx screens and random forest models, it was found that crRNA-folding energy, local target C and upstream target U context were the principal factors influencing knock-down efficacy in gRNA design (26). Consequently, various web-based tools emerged to facilitate Cas13 gRNA design. However, none has experimentally been assessed for its splicing editing efficacy. To test the capability of these new web-based tools to design efficient splice-editing gRNAs, we chose two well-established software for Cas13 gRNA design, CHOPCHOP (CC), which can be used to design gRNAs for several Cas13 family members (39), and Cas13design (CD), specific for CasRx (26). Both web tools calculate RNA accessibility and look for potential off-targets across the transcriptome for an increased specificity. Additionally, Cas13design considers gRNA-RNA hybridization energy, nucleotide preferences, crRNA folding and gRNA length (defined as 23nt by default) (26). We took advantage of each web tool to test top-scored gRNAs targeting *CTNND1* exon 2, in parallel of our strongest splice-editing custom-designed gRNA (Figure 1E-H). Both web tool-designed gRNAs, specially Cas13design gRNAs (gRNA_CD1 and CD2), showed stronger knock-down efficacies than our custom-designed gRNA_e2 (67% for CD1, 55% for CC1 and 42% for gRNA_e2, Figure 1H). However, at the splicing level, Cas13design gRNAs (specially gRNA_CD1) were more efficient than CHOPCHOP gRNAs, and comparable to our best splice-editing gRNA_e2 (Figure 1F), with no impact on overall gene expression levels (Figure 1G), turning this gRNA designer a suitable tool for dCasRx splice-editing. We next wanted to address how efficient the system is inducing a splice-switching effect at any alternatively spliced gene of interest.

### dCasRx is an efficient tool to induce accurate splice-switching effects at endogenously expressed transcripts

To evaluate dCasRx efficacy as a general tool to change splicing patterns at endogenously expressed genes, we targeted twelve additional exons from different genes that differ in exon length, transcript’s RNA abundance and basal splicing levels before editing, in case these were factors impacting dCasRx splice-editing efficiency (Figure 2A-D). Top-scored Cas13design gRNAs were preferentially used, although custom-designed gRNAs were also tested for regions in which no high-scored gRNA could be designed. Regardless of the exon length or transcript’s RNA abundance, dCasRx induced more than 2-fold changes in exon inclusion levels in 10/13 (77%) of the cases by just using a single gRNA targeting the exon of interest (Figure 2A-C and Supplementary Figure S2A-B). Even more, 70% of these splice-editing gRNAs induced a true splice-switching effect with changes in exon inclusion from included (PSI>40%) to excluded (PSI<15%) or with more than 40% changes in splicing (dPSI>40%) (Supplementary Figure S2A). However, despite targeting exons with already low inclusion levels before dCasRx targeting (PSI<10%, Figure 2D and Supplementary Figure 2B), most dCasRx-induced changes were towards further reduction in inclusion levels, with just one case of positive effect on *PKM* exon 9 (Figure 2A and Supplementary Figure S2A). This result is consistent with previous observations that dCasRx most likely regulate splicing by interfering with the splicing machinery (21, 26).

**Figure 2.**
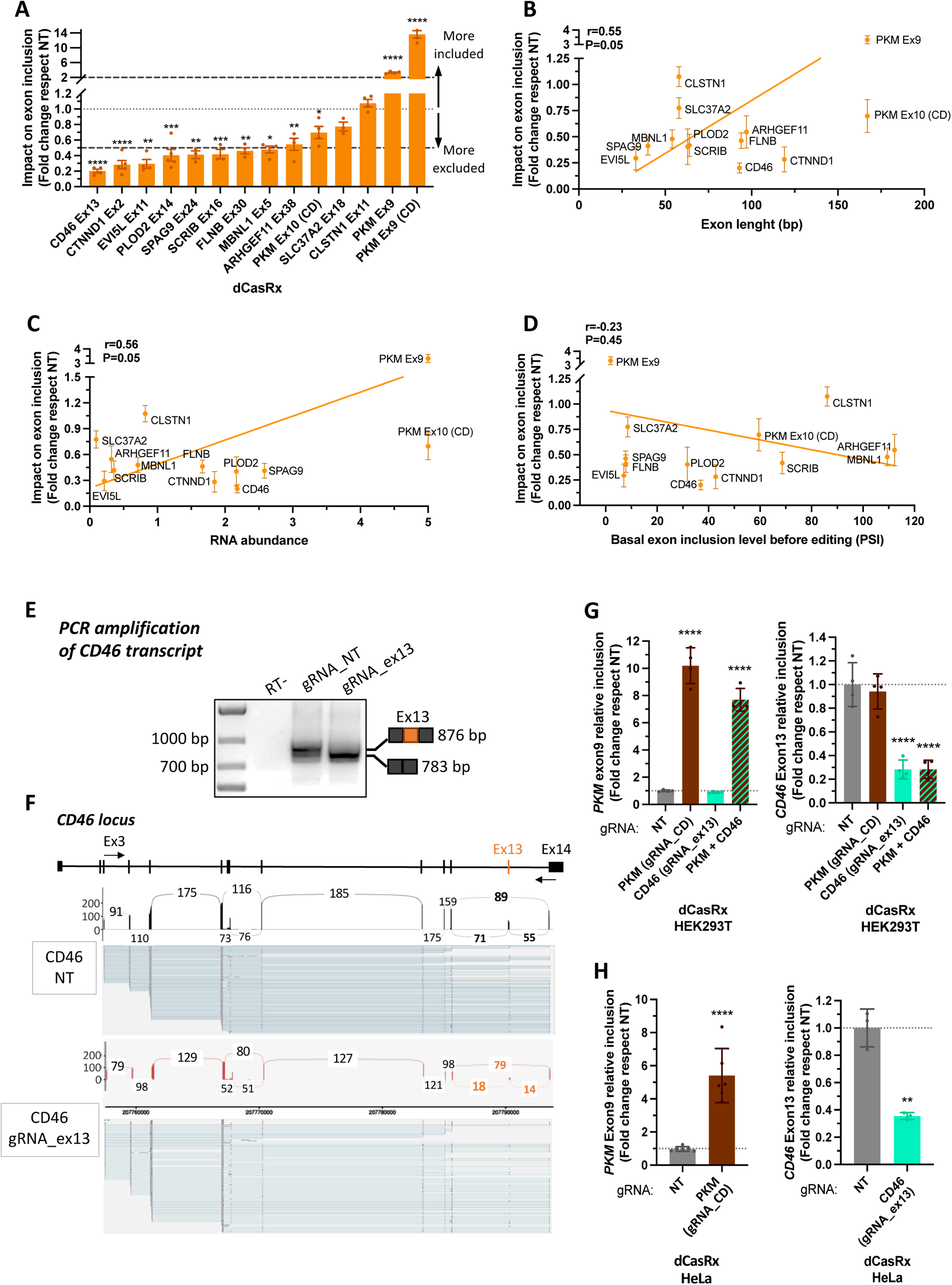
Efficient and accurate splice-editing of endogenous transcripts with dCasRx. (**A**) Changes in the inclusion levels of 13 exons targeted with dCasRx and their corresponding exon-specific gRNA in HEK293T cells. All exon inclusion levels were normalized to the values obtained with transfection of a non-targeting gRNA (NT) to evaluate the splicing editing efficiency as a fold change. (**B-D**) Pearson correlation between dCasRx splice-editing efficiency and the exons’ length (in bp) (B), total RNA abundance of the targeted transcripts (C) and basal inclusion levels (represented as PSI) of the targeted exons before dCasRx splicing editing in HEK293T cells (D). (**E**) Representative agarose gel showing *CD46* whole transcript (ex3 to ex14) after transfection of dCasRx and the strongest splice-editing gRNA. NT is used as a control. A shift in size is observed corresponding to changes in exon 13 inclusion. (**F**) Amplicon sequencing of the RT-qPCR shown in the agarose gel to visualize the changes in splicing across *CD46* upon targeting of dCasRx to exon 13. (**G**) *PKM* exon 9 and *CD46* exon 13 relative inclusion levels in HEK293T cells after transfection of dCasRx and either one or the two gRNAs inducing the strongest splice-editing effect. (**H**) *PKM* exon 9 and *CD46* exon 13 relative inclusion levels in HeLa cells after transfection of dCasRx and the strongest splice-editing gRNA. RT-qPCR levels were normalized by total expression of the corresponding gene. Data are represented as mean ± SD of the fold change relative to non-targeting gRNA (NT) in at least 3 biological replicates. *P <0.05, **P <0.01, ***P <0.001, **** P <0.0001 in unpaired T-tests (**A,H**) or unpaired one-way ANOVA (**G**) respect NT.

However, it has been proposed that fusing dCasRx to well-known splicing regulators, such as RBFOX1 or hnRNPA1, can improve splicing changes in either directions (21, 34). We thus compared the splice-switching efficacy of unconjugated dCasRx versus dCasRx fused to either RBFOX1 (Supplementary Figure 2C-F) or hnRNPA1 (Supplementary Figure 2G-I). Three exons differently impacted by dCasRx targeting were tested: *CD46* exon 13 and *PKM* exon 10 which splicing is reduced with different efficacities, and *PKM* exon 9 which is the only exon with increased inclusion (Figure 2A). Despite reducing *SMN2* exon 7 inclusion levels as previously reported (34), dCasRx-RBFOX1 did not reach the splicing changes obtained with dCasRx for none of the tested exons (Supplementary Figure S2C-F). On the other hand, dCasRx-hnRNPA1 had similar splicing effects as dCasRx (Supplementary Figure S2G-I), with a slight outperformance when targeted to *PKM* exon 10 (Supplementary Figure S2H). However, dCasRx-hnRNPA1 also induced a splicing effect at *PKM* exon 9 even in the absence of the targeting gRNA (NT condition), suggesting an indirect effect from overexpressing hnRNPA1’s effector’s domain (Supplementary Figure S2I). Altogether, our data support the use of unconjugated dCasRx combined with just one gRNA as a more specific splicing editor with less chances of off-target effects.

Another important factor to consider when inducing splice-switching effects by interfering with the splicing machinery is whether additional cryptic splice sites may also be activated, leading to the expression of novel isoforms. To evaluate this possibility we selected our best performing gRNA, targeting *CD46* exon 13, and amplified the whole transcript upon dCasRx targeting to sequence the amplicons with Oxford Nanopore long read sequencing (Figure 2E-F). Both agarose gel and amplicon sequencing confirmed a shift towards exclusion of exon 13 upon dCasRx targeting. Amplicon sequencing detected 41% of *CD46* transcripts including exon 13 in HEK293T cells (Figure 2F), which was also observed in the agarose gel with the presence of two bands corresponding to the transcripts including or not exon 13 (the exon is 93 bp, Figure 2E). Upon dCasRx targeting of exon 13, there was a clear shift in the agarose gel towards the lower band that reflected the exclusion of exon 13 in most of the transcripts, which was confirmed in the amplicon sequencing with a higher number of reads excluding exon 13 (79 reads vs 18/14 reads for inclusion). No cryptic splice sites were neither observed along *CD46* mRNA, supporting an accurate splice-switching effect of dCasRx specifically at the targeted region.

Finally, to further test dCasRx versatility and its potential to induce strong phenotypic changes, we targeted simultaneously two different exons and in a different cell line from HEK293T. dCasRx induced similar changes in *PKM* exon 9 and *CD46* exon 13 splicing, whether targeted individually or simultaneously at both exons (Figure 2G). Furthermore, comparable splicing effects were obtained in epithelial HeLa cells (Figure 2H), increasing the interest of this new RNA tool for studying splicing in any cellular context.

In conclusion, unconjugated dCasRx is an efficient, easy to implement and cost-effective tool to induce specific splice-switching changes in the alternative splicing of endogenous genes. Furthermore, we have identified a web-based software dedicated to Cas13 targeting, called Cas13design, that is also suitable for designing splice-editing gRNAs. However, as already mentioned with *CTNND1* exon 2, we noticed that despite targeting an exon with different gRNAs successfully degrading the RNA when targeting CasRx (Figure 1D), there were not all as efficient editing splicing (Figure 1B), suggesting a position-dependent functional effect. Differences in secondary and tertiary RNA structures between CasRx-targeting spliced mRNAs and dCasRx-targeting unspliced pre-mRNAs could explain these differences in splice-editing capacity between gRNAs. Of note, a similar position effect has been reported when targeting dCasRx with 32 tiling gRNAs to a poison exon in *TRA2β*, supporting a biologically meaningful position-dependent effect of dCasRx in splicing (4).

### dCasRx position-dependent splicing effect reflects the targeting of key cis-regulatory regions along the alternatively spliced exon

To evaluate the aforementioned position-dependent splicing effect, we designed 3 to 4 tiling gRNAs across four additional alternatively spliced exons that cover the whole exonic region (Figure 3A,E and Supplementary Figure S3A,E). Again, despite successfully inducing an RNA knock-down with most of the designed gRNAs (Figure 3D,H and Supplementary Figure S3D,H), only specific gRNAs induced a splice-switching effect higher than two-fold when targeting dCasRx to the tested exons (Figure 3B,F and Supplementary Figure S3B,F). Even more, there was no correlation between CasRx knock-down and dCasRx splice-editing efficiency, despite using the same targeting gRNAs with both systems (Supplementary Figure S3I). These results suggest that the observed position-dependent splicing effect extends beyond the gRNA’s capability to target the RNA. Instead, we hypothesize that it might underline the targeting of key cis-regulatory elements influencing alternative splicing.

**Figure 3.**
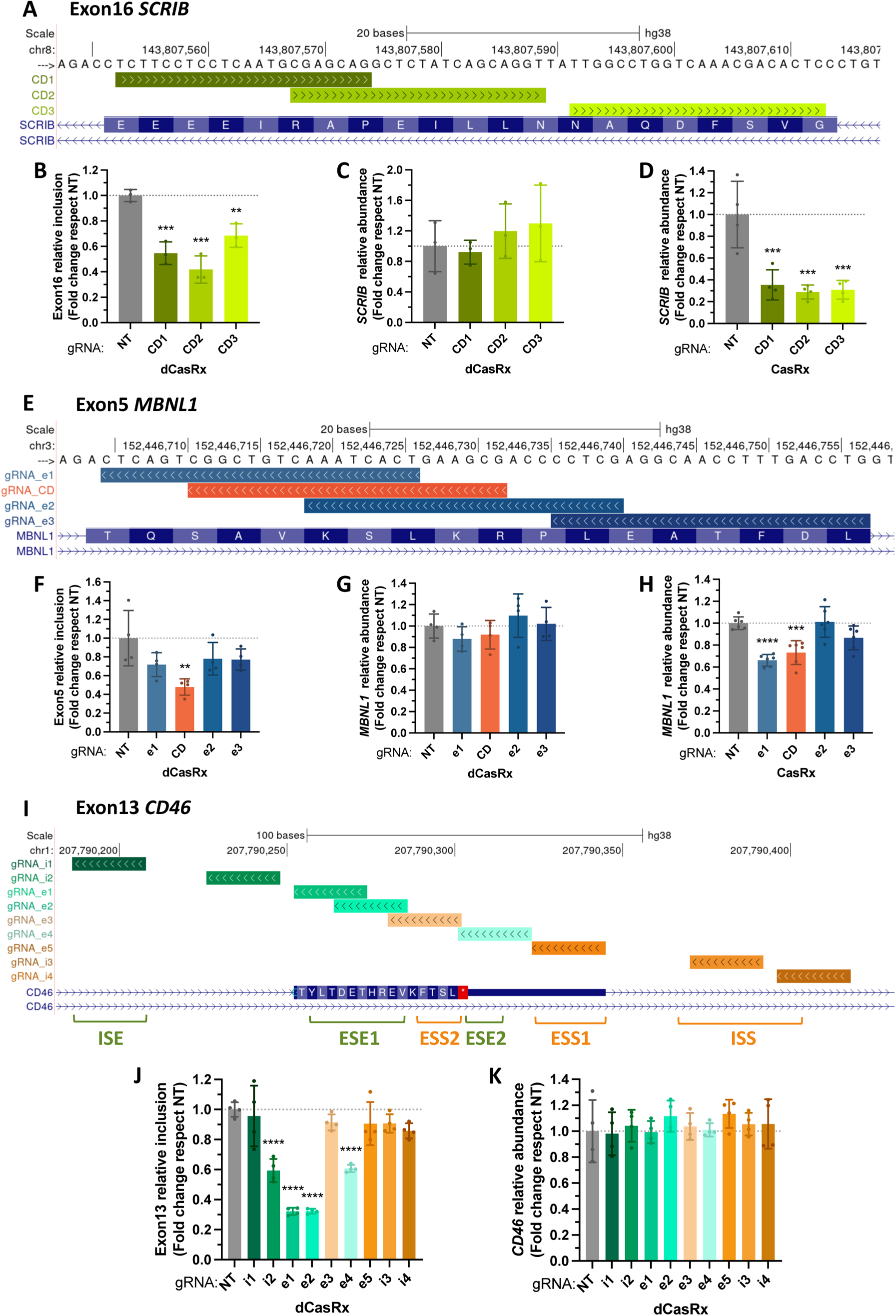
dCasRx position-dependent splicing effect underlines the existence of key cis-regulatory RNA elements at the targeted exon. (**A**) Genomic location of gRNAs tiled across *SCRIB* exon 16. (**B**) *SCRIB* exon 16 relative inclusion levels in HEK293T cells after transfection of dCasRx and the indicated gRNAs. (**C,D**) *SCRIB* relative RNA abundance after transfection of dCasRx (C) or CasRx (D). (**E**) Genomic location of gRNAs tiled across *MBNL1* exon 5. (**F**) *MBNL1* exon 5 relative inclusion levels after transfection of dCasRx and the indicated gRNAs. (**G,H**) *MBNL1* relative RNA abundance after transfection of dCasRx (G) or CasRx (H). (**I**) Genomic location of 9 gRNAs tiled across *CD46* exon 13 and two flanking intronic regions reported to regulate its splicing (ISE and ISS). (**J**) *CD46* exon 13 relative inclusion levels after transfection of dCasRx and the indicated gRNAs. (**K**) *CD46* relative RNA abundance after transfection of dCasRx and the indicated gRNAs. RT-qPCR levels were normalized by total expression levels of the corresponding gene for splicing analysis; while RNA abundance was normalized to *TBP* housekeeping gene expression. Data are represented as mean ± SD of the fold change relative to non-targeting gRNA (NT) in at least 3 biological replicates. *P <0.05, **P <0.01, ***P <0.001, **** P <0.0001 in unpaired one-way ANOVA respect NT.

To evaluate this hypothesis, we targeted an exon with well-known regulatory elements in a gene important for the immune system, *CD46* (40). Thanks to an extensive characterization in splicing reporters (40), *CD46* exon 13 is known to be regulated by two strong exonic splicing enhancers (ESE), two exonic silencers (ESS) and several intronic silencers (ISS) and enhancers (ISE) (Figure 3I). As expected, targeting any of the two exonic splicing enhancers (ESE1 or ESE2) strongly reduced *CD46* exon13 inclusion, without impacting overall gene expression levels (Figure 3J-K). While targeting the exonic splicing silencers (ESS2 or ESS1) did not have such effect. Surprisingly, none of the gRNAs targeting the identified intronic splicing enhancer (ISE) or silencer (ISS) had an effect on *CD46* splicing, except for gRNA_i2 that is overlapping the branch point and slightly the acceptor site. This is consistent with previous observations in which targeting dCasRx to intronic regions flanking an alternatively spliced exon, like *TRA2*β or *SMN2*, had a milder effect than targeting exonic regions (4, 34). Finally, if such position-dependent effect was indeed the result of targeting specific regulatory RNA elements across the exon, targeting of constitutive exons, whose splicing is more robust and less enriched in these intermediate regulatory sequences (41), should be more difficult to impact. None of the gRNAs tiled across three constitutive exons present at *CD46* or *PKM* transcripts impacted splicing (Supplementary Figure S3K-N, S3P-S), even though the designed gRNAs were capable of targeting CasRx for RNA degradation (Supplementary Figure S3O,T).

These results support our hypothesis that dCasRx position-dependent effect is the reflection of the targeting of key cis-regulatory sequences along the pre-mRNA, with a preference for exonic over intronic regions.

### A novel application for dCasRx to identify mechanisms of alternative splicing regulation

If dCasRx regulates splicing by targeting regulatory regions, it is most likely doing so by interfering with the recruitment of key splicing factors to these regions. We thus next targeted an alternatively spliced gene whose splicing regulators are well-known, the Muscle Pyruvate Kinase (*PKM*). *PKM* is a glycolytic enzyme that is spliced into two mutually exclusive isoforms. *PKM1* includes exon 9 and is expressed in non-proliferative cells. While the switch to exon 10 leads to *PKM2* isoform that confers a pro-oncogenic metabolic advantage such that cancer cells can take up more glucose than normal tissues (42, 43). Targeting dCasRx to *PKM* exon 9 (Figure 4A) or exon 10 (Figure 4E) with tiling gRNAs across the two exons highlighted once again clear differences in their mechanisms of splicing regulation. The strongest splice-editing gRNA targeting exon 9 (gRNA_e5) is actually increasing the exon’s inclusion (Figure 4B and Supplementary Figure S4A). This gRNA is complementary to two well-known RNA motifs for the splicing repressor hnRNPA1, which are known to inhibit exon 9 inclusion (45) (Figure 4A). If dCasRx was interfering with the spliceosome, it should have decreased exon 9 inclusion. Instead, we observed an increase similar to the one observed upon knock-down of hnRNPA1 (Figure 4D and Supplementary Figure S4D), which is consistent with dCasRx interfering with hnRNPA1 recruitment to its RNA binding sites along the pre-mRNA.

**Figure 4.**
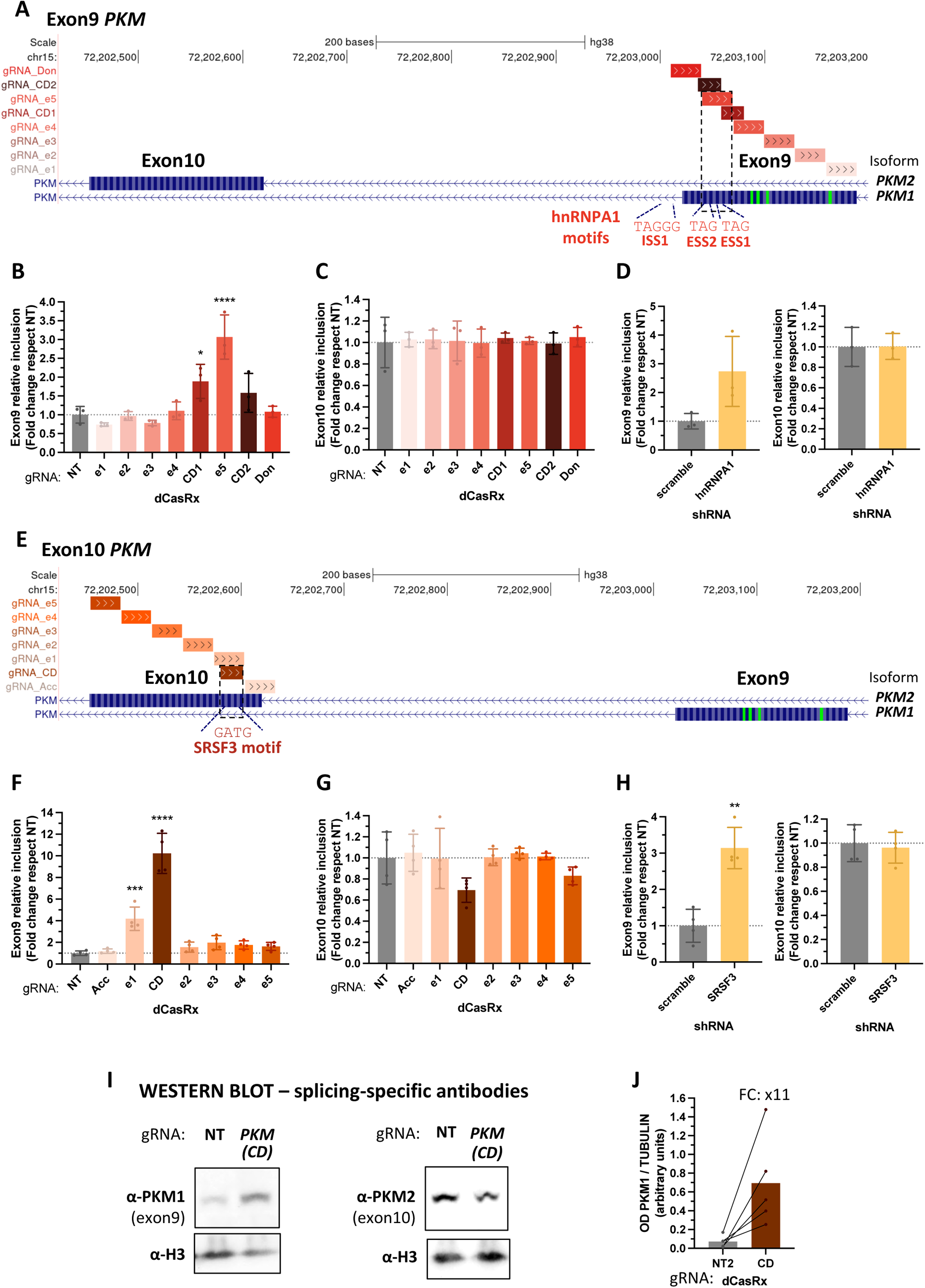
A novel role for dCasRx to identify mechanisms of alternative splicing regulation. (**A,E**) Genomic location of gRNAs tiled across *PKM* exon 9 (A) or exon 10 (E). hnRNPA1 and SRSF3 RNA binding motifs are shown. (**B-C, F-G**) *PKM* exon 9 and *PKM* exon 10 relative inclusion levels in HEK293T cells after transfection with dCasRx and the indicated gRNAs. (**D,H**) *PKM* exon 9 and exon 10 relative inclusion levels in HEK293T cells transfected with shRNAs against hnRNPA1 (D) or SRSF3 (H). (**I**) Representative Western Blot detecting specifically PKM1 (includes exon 9) or PKM2 (includes exon 10) isoforms after transfection of dCasRx and the strongest splice-editing gRNA. TUBULIN and H3 proteins are used as loading controls. (**J**) Quantification of PKM1 Western blots bands with ImageJ software. The optical density (OD) of each band was normalized by the corresponding TUBULIN’s OD for each of the 5 biological replicates. Exons 9 and 10 RT-qPCR levels were normalized by total *PKM* expression levels. Data are represented as mean ± SD of the fold change relative to non-targeting gRNA (NT) in at least 4 biological replicates. *P <0.05, **P <0.01, ***P <0.001, **** P <0.0001 in unpaired one-way ANOVA (**B-C,F-G**) respect NT or T-test (**D,H**) respect scramble shRNA

Even more, highlighting the capacity of dCasRx to unveil novel mechanisms of splicing regulation, we found that the gRNA with the strongest splice-switching effect on *PKM* exon 9, both at the RNA and protein level, is in fact targeting exon 10 (gRNA_CD, Figure 4E-G, I-J and Supplementary Figure S4E-G, SI-J). Indeed, this gRNA not only decreased exon 10 inclusion levels of 30%, but it also induced a 10x-fold increase in the inclusion levels of the mutually exclusive exon 9, from 2% to 23% (Supplementary Figure S4E-F), that was translated into an 11x-fold increase in PKM1 (includes exon9) protein isoform levels respect NT (Figure 4I-J and Supplementary Figure S4I-J). Amplicon sequencing of full-length *PKM* cDNA confirmed the correct switch from exon 10 to exon 9 inclusion in 27% of the transcripts upon dCasRx targeting (Supplementary Figure S4K-L), supporting again an accurate splice-switching effect of dCasRx that reaches the protein level.

When looking for RNA motifs along the 23-bp region targeted by this splice-switching gRNA (Figure 4E), we identified SRSF3, which has previously been reported to inhibit exon 9 inclusion by binding to exon 10 (43, 44, 46). In line with our results, SRSF3 knock-down increased exon 9 inclusion (Figure 4H and Supplementary Figure S4H). Whereas SRSF3 overexpression reduced of 35% dCasRx effect on exon 9 splicing (Figure 5A-B and Supplementary Figure S5A-B). To confirm that the dCasRx effect on *PKM* exon 9 inclusion is by competing with SRSF3 recruitment to *PKM* pre-mRNA, we performed an RNA immunoprecipitation assay. As expected, SRSF3 binding to *PKM* exon 10 pre-mRNA decreased by 35% in the presence of the targeting gRNA_CD, with no impact on SRSF3 binding to its own pre-mRNA, used as a positive control (Figure 5D and Supplementary Figure S5C).

**Figure 5.**
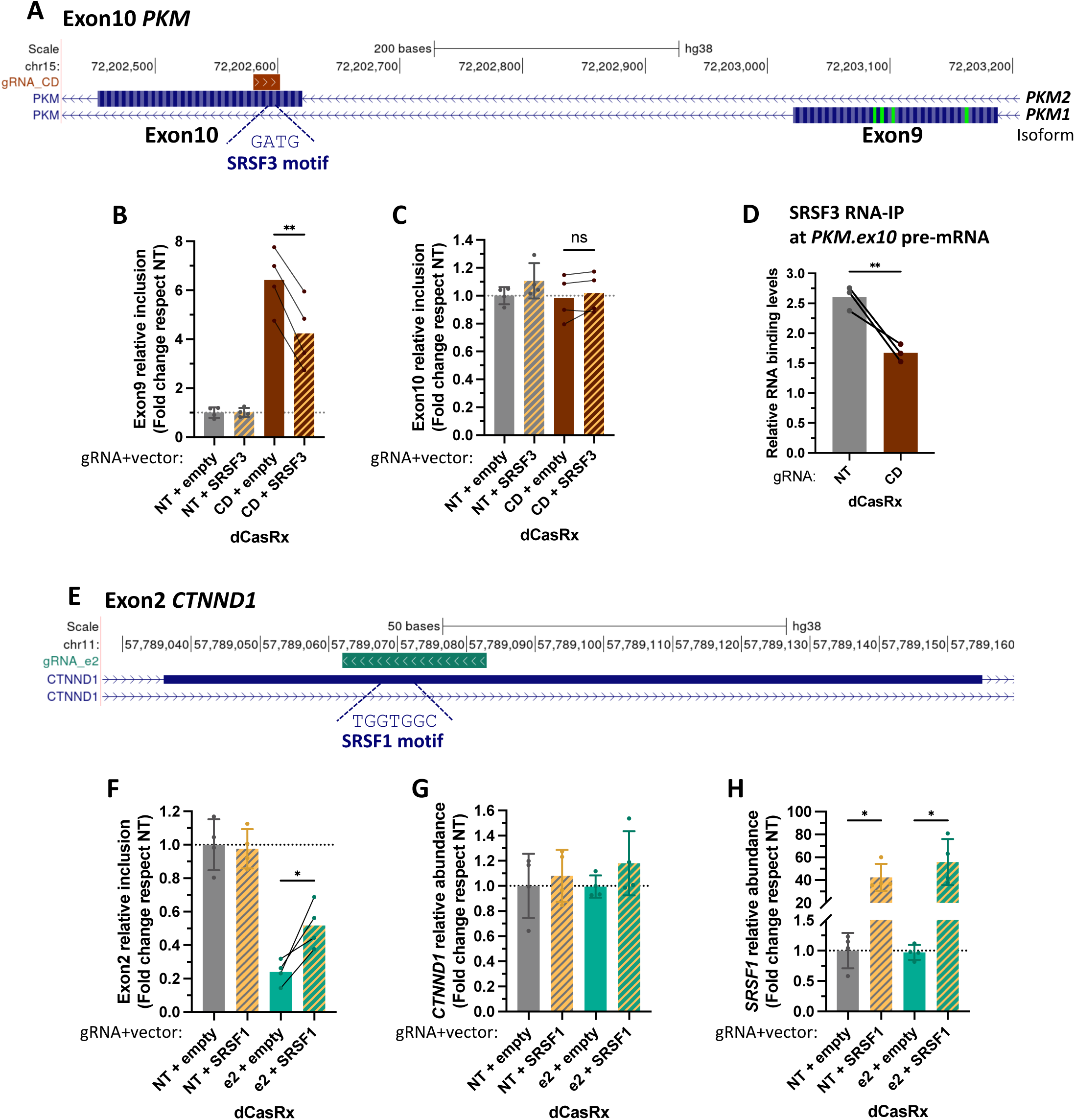
dCasRx interferes with recruitment of the splicing factors to the pre-mRNA. (**A**) Genomic location of *PKM*’s strongest splice-editing gRNA and SRSF3 RNA motif at exon 10. (**B-C**) *PKM* exon 9 and exon 10 relative inclusion levels in HEK293T cells transfected with dCasRx and the splice-editing gRNA_CD upon SRSF3 overexpression. Non-targeting gRNA (NT) and empty vector were used as controls. (**D**) SRSF3 recruitment levels to *PKM* pre-mRNA measured by RNA immunoprecipitation (RNA-IP) upon transfection of dCasRx and the splice-editing gRNA_CD. SRSF3 immunoprecipitation levels at *PKM* exon 10 pre-mRNA were normalized by the recruitment levels obtained at the positive control pre-mRNA in 3 biological replicates. (**E**) Genomic location of *CTNND1*’s strongest splice-editing gRNA and SRSF1 predicted RNA motif at exon 2. (**F**) *CTNND1* exon 2 relative inclusion levels in HEK293T cells transfected with dCasRx and the splice-editing gRNA_e2 upon SRSF1 overexpression. NT gRNA and empty vector were used as controls (**G-H**) *CTNND1* and *SRSF1* relative RNA levels. RT-qPCR levels were normalized by *PKM* or *CTNND1* total expression levels for splicing analysis; while RNA abundance was normalized by *TBP* housekeeping gene expression. In **B-C** and **F-H**, data are represented as mean ± SD of the fold change relative to cells transfected with non-targeting gRNA (NT) and empty vector in at least 4 biological replicates. *P <0.05, **P <0.01, ***P <0.001, **** P <0.0001 in paired T-test respect empty (**B,C,F,G,H**) or unpaired T-Test respect NT (**D**).

Finally, to test the new role of dCasRx in identifying novel splicing regulators, we investigated *CTNND1* splicing, which regulators are not entirely known yet. RNA motif search analysis combined with shRNA knock-down of the predicted splicing factors along *CTNND1* exon 2, pointed to SRSF1 as the strongest activator of exon 2 splicing (Supplementary Figure S5D-F). Since the gRNA inducing the most efficient splice-switching effect on *CTNND1* is precisely targeting this SRSF1 binding site (gRNA_e2, Figure 5E and S5D), we tested the effect of overexpressing SRSF1 on dCasRx editing. In concordance with a competition effect with the splicing activator, dCasRx was not capable of inhibiting exon 2 inclusion efficiently in the presence of SRSF1 (Figure 5F-H). Altogether, these results validate the role of dCasRx in pointing out succinct RNA regions to identify key regulators of alternative splicing without the need of downregulating a battery of potential splicing factors predicted from the search analysis of degenerated RNA motifs along the entire exon.

## CONCLUSION

We propose dCasRx as a versatile, cost-effective and easy-to-implement RNA tool to switch the alternative splicing of endogenously expressed genes, both at the RNA and protein level, for proper study of the role of a specific splicing event in its physiological context. Furthermore, efficient splice-editing effects are obtained with just one gRNA per exon and unconjugated dCasRx, with no need to fuse the dCasRx to a splicing factor domain, contrary to what was previously reported (21, 34), which dramatically reduces potential off-target effects.

We have also observed that accessibility to the RNA is not the only factor important for the designing of strong splice-editing gRNAs. Splice-switching effects were obtained when targeting key regulatory RNA elements along the alternatively spliced exon. We thus encourage to test several gRNAs tiled across the exon of interest for selection of the strongest splice-editing gRNA. Furthermore, with such strategy, key regulatory RNA elements will also be identified, as a new application of the dCasRx system that does not require complicated experimental settings. Effectively, splicing enhancer and silencer domains, together with specific splicing factor binding sites, have classically been identified by deletion or mutation of putative RNA sequences in mostly splicing reporter minigenes (4, 40, 45). Unfortunately, such systems do not often reflect the complexity of an endogenously expressed gene, which is surrounded by a specific chromatin environment that can impact co-transcriptional splicing (47), and in which the exon is flanked by longer intronic regions prone to form complex secondary structures, also impacting splicing. Using competition assays in two different alternatively spliced exons at *PKM* and *CTNND1* transcripts, we have shown that dCasRx can also be used to identify key regulatory binding sites without the need of complex deletion/mutants. As a caveat, dCasRx seems to target more efficiently exonic than intronic regulatory regions, which can be a limiting factor when competing against splicing repressors that often bind to intronic RNA elements and at more than one site around the regulated exon (48). Since we have observed that dCasRx can efficiently induce splicing changes at two different sites at the same time using two different gRNAs, we propose to expand the tiling of gRNAs at flanking introns, but combining multiple targeting gRNAs for improved repressor effects to induce exon inclusion and identify like this key intronic splicing regulators.

Finally, CRISPR-Cas9 DNA editing systems have successfully corrected splicing-related mutations *in vivo* (49). However, the risk of permanent genome editing and secondary off-target sites questions its use in the clinic. Since dCasRx targets the pre-mRNA, it can rescue disease-associated splice variants with no need to introduce a permanent change in the genome. Furthermore, dCasRx not only efficiently shifts alternative splicing outcome into the correct splicing isoform, evidenced by amplicon sequencing of the edited transcripts, but it can do so to more than one alternatively spliced event by co-transfecting multiple gRNAs. This will significantly increase the phenotypic impact when correcting complex splicing-associated anomalies in which more than one splicing isoform is implicated, as in cancer or brain diseases (50–52). However, as a second caveat, we encountered some difficulties reaching functionally-relevant dCasRx expression levels in difficult-to-transfect cells. The use of programable Cas13 viral particles to edit primary cells, such as neurons, or to correct disease phenotypes *in vivo* with inducible CasRx, are proving to be valuable delivering tools for safer CRISPR-based therapeutic strategies (24, 26, 53, 54). We thus propose CRISPR-dCasRx as a versatile RNA-targeting tool to study alternative splicing from a mechanistic and biological point of view.

## ACKNOWLEDGEMENTS

We thank Alvaro Herrero-Reiriz for his support in the analysis of the amplicon sequencing data. Dr. Maria Moriel-Carretero for her very insightful comments about the manuscript. Dr. Laguette (IGH, Montpellier) and Dr. Amor-Gueret (Institut Curie, Orsay) for HEK293T and HeLa cell lines, respectively. Dr. Patrick Hsu (Salk, US) for providing us the plasmids of CasRx, dCasRx and processed gRNA through Addgene. We thank Dr. Ling-Ling-Ling Chen Lab (State Key Laboratory of Molecular Biology, Shanghai Institute of Biochemistry and Cell Biology, China) for providing us the plasmids for PspCas13b, dPspCas13b, dPguCas13b and crRNA backbone. We thank Dr. Andriana Margariti (Queen’s University Belfast, Ireland) for SRSF3 shRNA, and Dr. Olga Anczuków (Jackson Laboratory, USA) for SRSF1 and SRSF3 mammalian expression vectors.

