## Supplemental Figures for "“A CRISPR-dCas13 RNA-editing tool to study alternative splicing”"

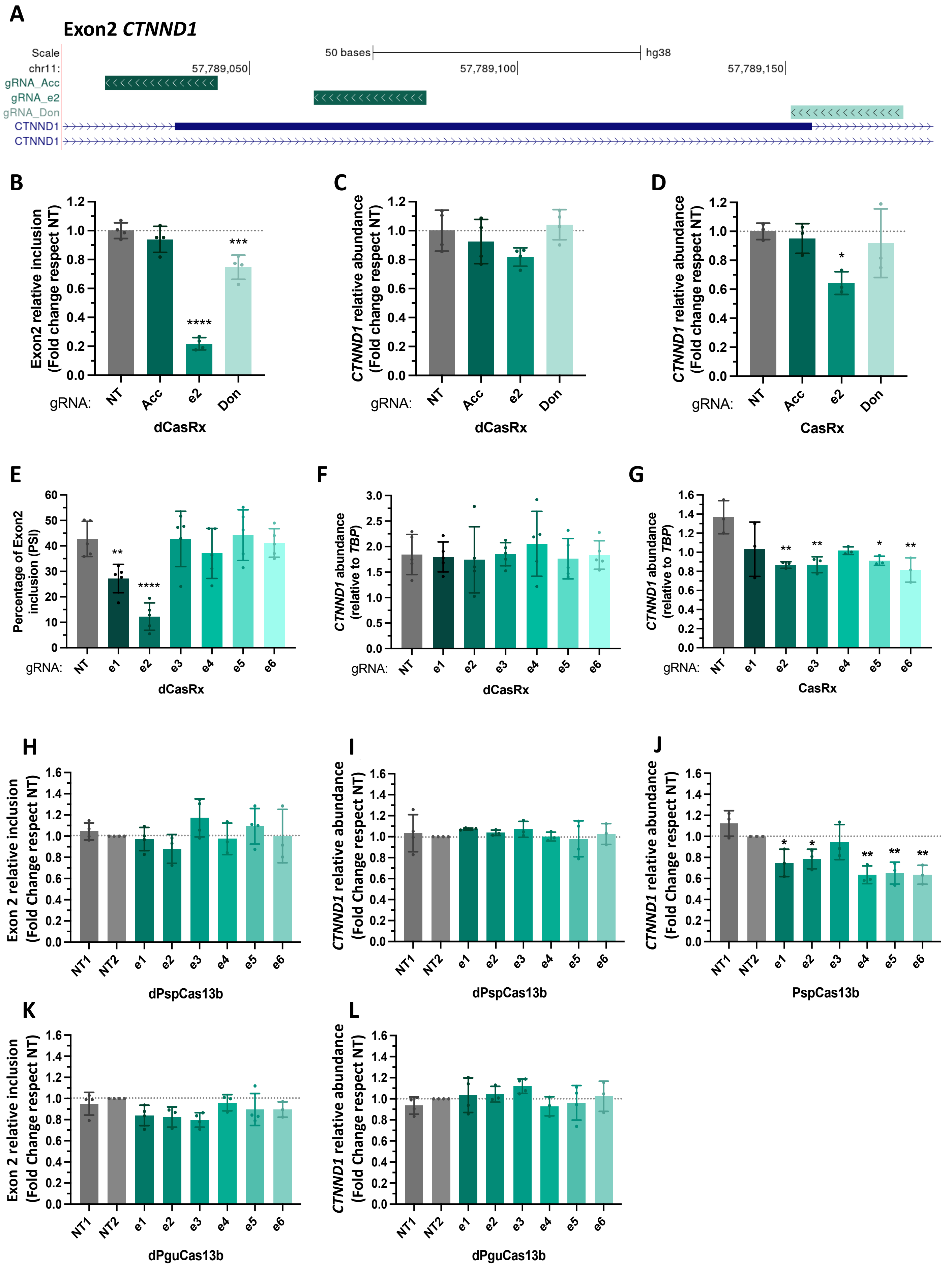
A
Exon2 CTNND1
Scale
50 bases
hg38
chr11:
57,789,050
57,789,100
57,789,150
gRNA_Acc
gRNA_e2
gRNA_Don
CTNND1
CTNND1
B
Exon2 relative inclusion (Fold change respect NT)
gRNA: NT Acc e2 Don
dCasRx
C
CTNND1 relative abundance (Fold change respect NT)
dCasRx
D
CTNND1 relative abundance (Fold change respect NT)
CasRx
E
Percentage of Exon2 inclusion (PSI)
gRNA: NT e1 e2 e3 e4 e5 e6
dCasRx
F
CTNND1 abundance (relative to TBP)
dCasRx
G
CTNND1 abundance (relative to TBP)
CasRx
H
Exon 2 relative inclusion (Fold Change respect NT)
NT1 NT2 e1 e2 e3 e4 e5 e6
dPspCas13b
I
CTNND1 relative abundance (Fold Change respect NT)
dPspCas13b
J
CTNND1 relative abundance (Fold Change respect NT)
PspCas13b
K
Exon 2 relative inclusion (Fold Change respect NT)
dPguCas13b
L
CTNND1 relative abundance (Fold Change respect NT)
dPguCas13b

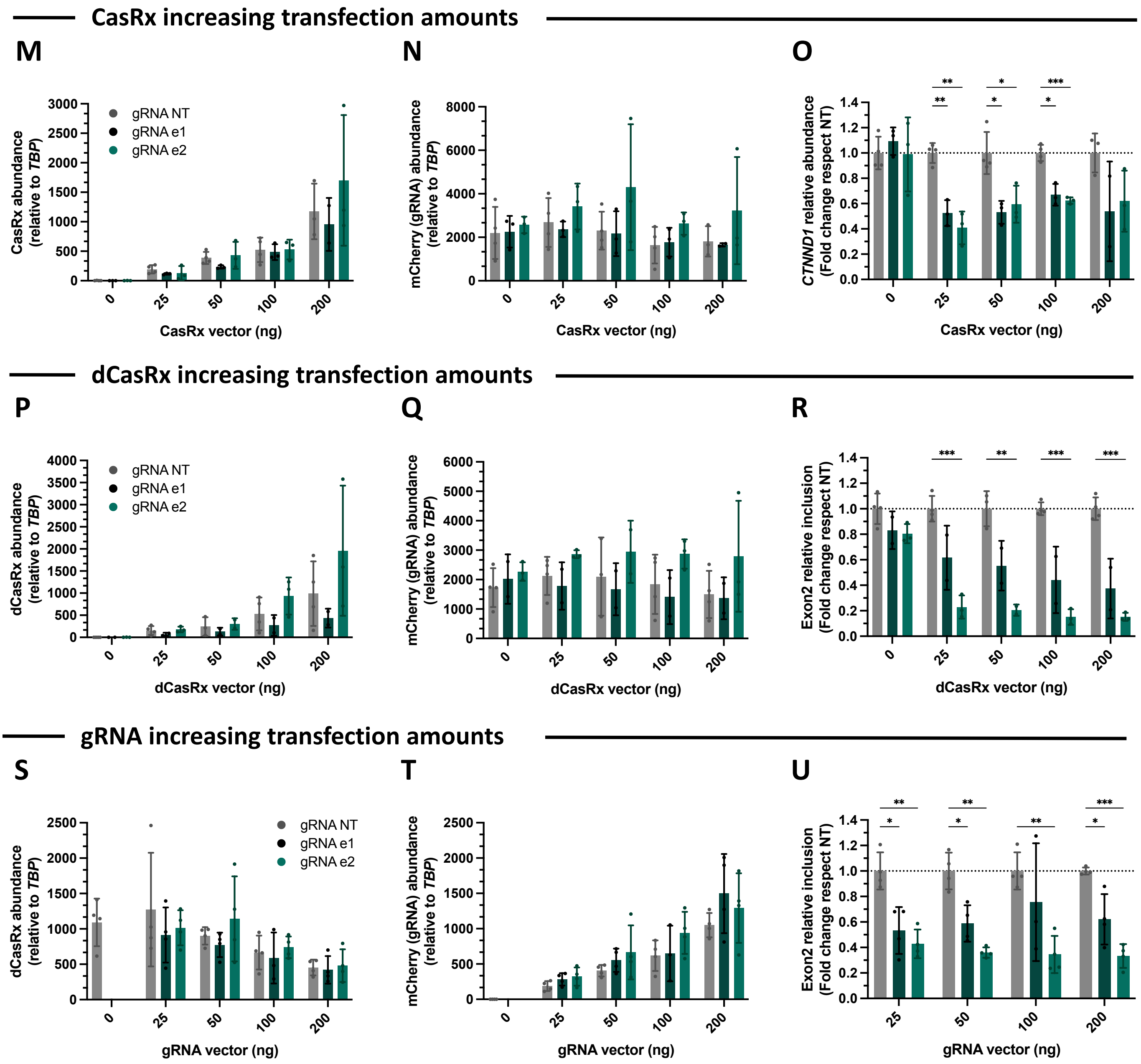


**Supplementary Figure 1. dCasRx is the dCas13 of choice for editing splicing.** (**A**) Genomic location of the acceptor and donor sites at *CTNND1* exon 2. As a reference we kept the strongest splice-editing gRNA_e2. (**B**) *CTNND1* exon 2 inclusion levels relative to non-targeting (NT) gRNA in HEK293T cells after transfection of dCasRx and the indicated gRNAs. (**C, D**) *CTNND1* RNA abundance relative to NT after transfection of dCasRx (**C**) or CasRx (**D**) and the indicated gRNAs. (**E**) Percentage of *CTNND1* exon 2 inclusion after transfection of dCasRx and the indicated gRNAs. (**F,G**) *CTNND1* RNA abundance after transfection of dCasRx (**F**) or CasRx (**G**) and the indicated gRNAs. (**H,K**) *CTNND1* exon2 inclusion levels relative to NT after transfection of the catalytically inactive dPspCas13b (**H**) or dPguCas13b (**K**) and the indicated gRNAs. (**I,J,L**) *CTNND1* RNA abundance relative to NT after transfection of dPspCas13b (**I**), dPguCas13b (**L**) or the catalytically active PspCas13b (**J**) and the indicated gRNAs. (**M-O**) CasRx, gRNA expression levels (assessed by expression of an mCherry reporter in frame with the gRNA) and *CTNND1* RNA abundance relative to NT in HEK293T cells transfected with 200 ng of the indicated gRNA and increasing amounts of CasRx (0-200ng). (**P-R**) dCasRx, gRNA expression levels (assessed by expression of an mCherry reporter in frame with the gRNA) and *CTNND1* exon 2 inclusion levels relative to NT in HEK293T cells transfected with 200 ng of the indicated gRNA and increasing amounts of dCasRx (0-200ng). (**S-U**) dCasRx, gRNA expression levels (assessed by expression of an mCherry reporter in frame with the gRNA) and *CTNND1* exon 2 inclusion levels relative to NT in HEK293T cells transfected with 200 ng of dCasRx and increasing amounts of the indicated gRNA (0-200ng). Exon 2 RT-qPCR levels were normalized by *CTNND1* total expression levels; while CasRx, dCasRx, mCherry and *CTNND1* total expression levels were normalized by housekeeping *TBP*. Data are represented as mean ± SD in at least 3 biological replicates. *P <0.05, **P <0.01, ***P <0.001, **** P <0.0001 in unpaired (**B-D, H-L, M-U**) or paired (**E-G**) one-way ANOVA respect NT.

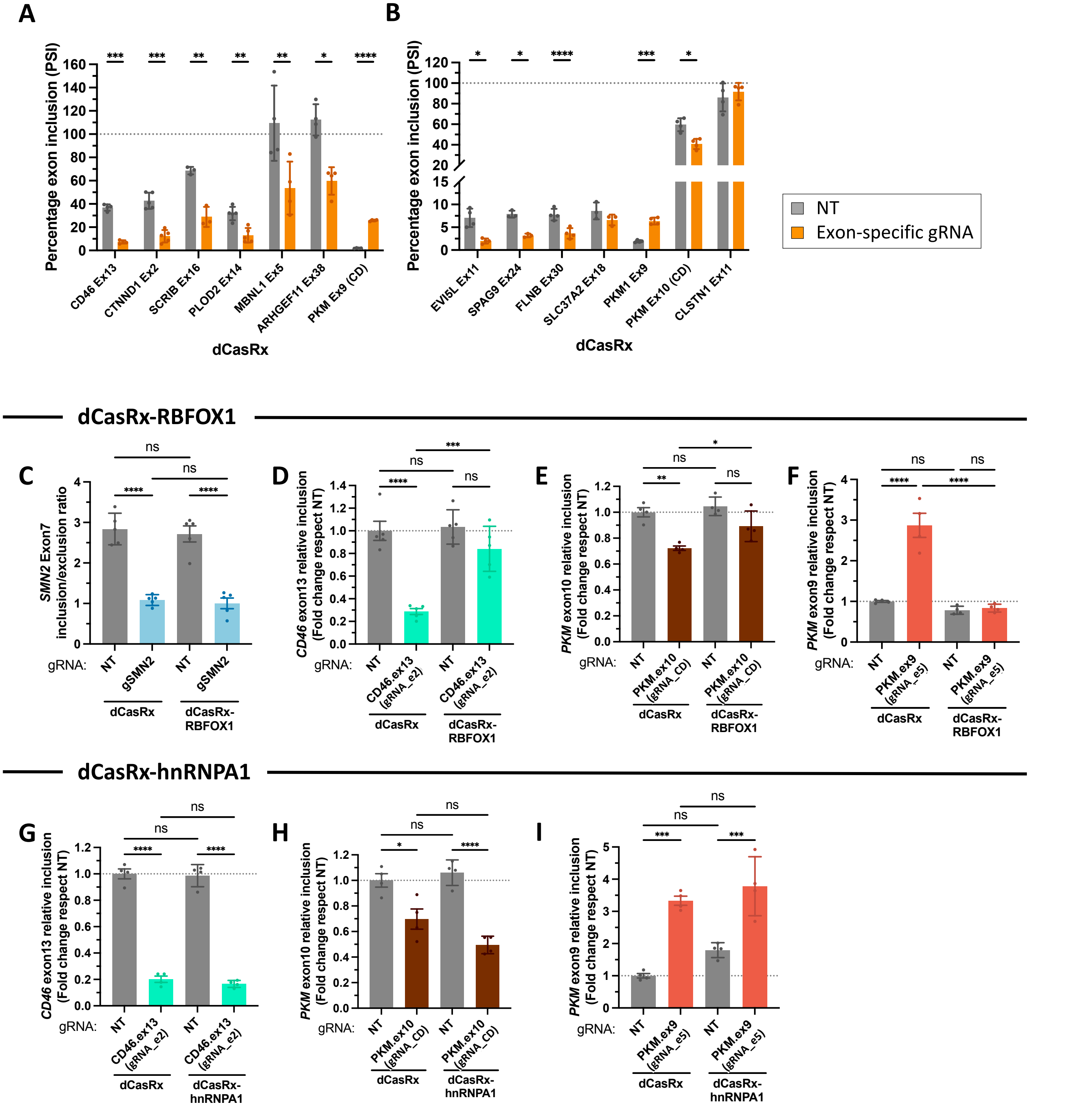


**Supplementary Figure 2. dCasRx alone can efficiently induce splice-switching of endogenous transcripts (A-B)** Percentage of exon inclusion (PSI) at 13 endogenously expressed transcripts in HEK293T cells transfected with dCasRx and either a gRNA targeting the studied alternatively spliced exon (in orange) or a non-targeting gRNA (NT, in grey). (**C-I**) Splicing editing efficiency of dCasRx compared to dCasRx-RBFOX1 (C-F) or dCasRx-hnRNPA1 (G-I) targeted to *SMN2* exon 7 (C), *CD46* exon 13 (D,G), *PKM* exon 10 (E,H) or *PKM* exon 9 (F,I). All exon inclusion values were normalized to the values obtained upon transfection with dCasRx and NT gRNA. RT-qPCR levels were normalized by total gene expression of the corresponding transcript to evaluate the percentage of inclusion (PSI). Data were represented as mean ± SD of the percentage of exon inclusion (**A-B**) or a fold change relative to non-targeting gRNA (NT) (**C-I**) in at least 4 biological replicates. *P <0.05, **P <0.01, ***P <0.001, **** P <0.0001 in paired T-tests respect NT (**A-B**) or unpaired one-way ANOVA (**C-I**).

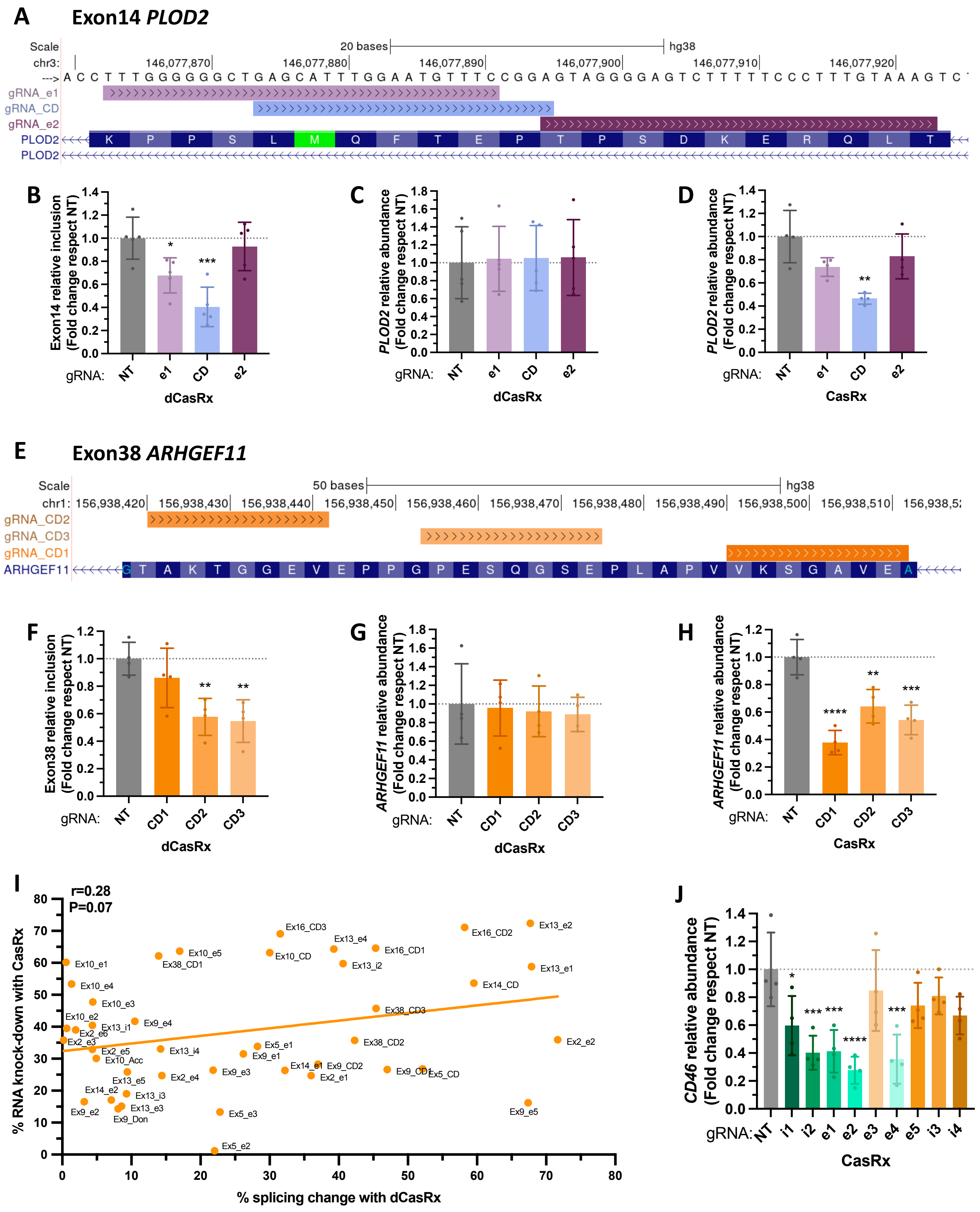
A
Exon14 PLOD2
Scale
20 bases
hg38
chr3:
146,077,870
146,077,880
146,077,890
146,077,900
146,077,910
146,077,920
gRNA_e1
gRNA_CD
gRNA_e2
PLOD2
PLOD2
B
Exon14 relative inclusion (Fold change respect NT)
gRNA: NT e1 CD e2
dCasRx
C
PLOD2 relative abundance (Fold change respect NT)
gRNA: NT e1 CD e2
dCasRx
D
PLOD2 relative abundance (Fold change respect NT)
gRNA: NT e1 CD e2
CasRx
E
Exon38 ARHGEF11
Scale
50 bases
hg38
chr1: 156,938,420
156,938,430
156,938,440
156,938,450
156,938,460
156,938,470
156,938,480
156,938,490
156,938,500
156,938,510
gRNA_CD2
gRNA_CD3
gRNA_CD1
ARHGEF11
F
Exon38 relative inclusion (Fold change respect NT)
gRNA: NT CD1 CD2 CD3
dCasRx
G
ARHGEF11 relative abundance (Fold change respect NT)
gRNA: NT CD1 CD2 CD3
dCasRx
H
ARHGEF11 relative abundance (Fold change respect NT)
gRNA: NT CD1 CD2 CD3
CasRx
I
r=0.28
P=0.07
% RNA knock-down with CasRx
% splicing change with dCasRx
Ex16_CD3
Ex13_e4
Ex16_CD1
Ex16_CD2
Ex13_e2
Ex10_e1
Ex10_e5
Ex38_CD1
Ex10_CD
Ex13_i2
Ex13_e1
Ex10_e4
Ex14_CD
Ex10_e3
Ex38_CD3
Ex10_e2
Ex13_i1
Ex9_e4
Ex2_e6
Ex2_e3
Ex5_e1
Ex38_CD2
Ex2_e2
Ex2_e5
Ex13_i4
Ex9_e1
Ex10_Acc
Ex14_e1
Ex9_CD2
Ex13_e5
Ex2_e4
Ex9_e3
Ex2_e1
Ex9_CD1
Ex5_CD
Ex14_e2
Ex13_i3
Ex13_e3
Ex9_e2
Ex9_Don
Ex5_e3
Ex9_e5
Ex5_e2
J
CD46 relative abundance (Fold change respect NT)
gRNA: NT i1 i2 e1 e2 e3 e4 e5 i3 i4
CasRx

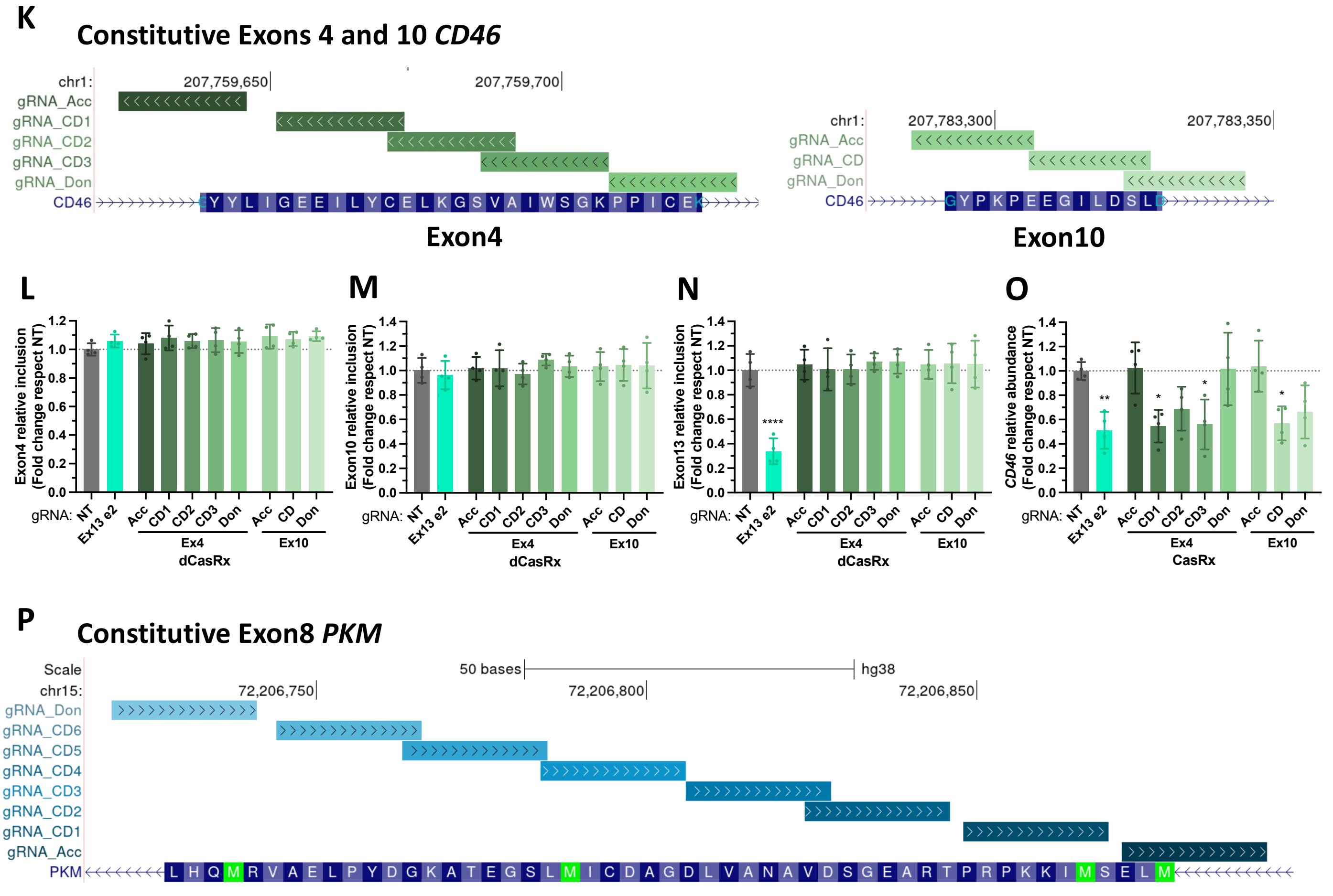


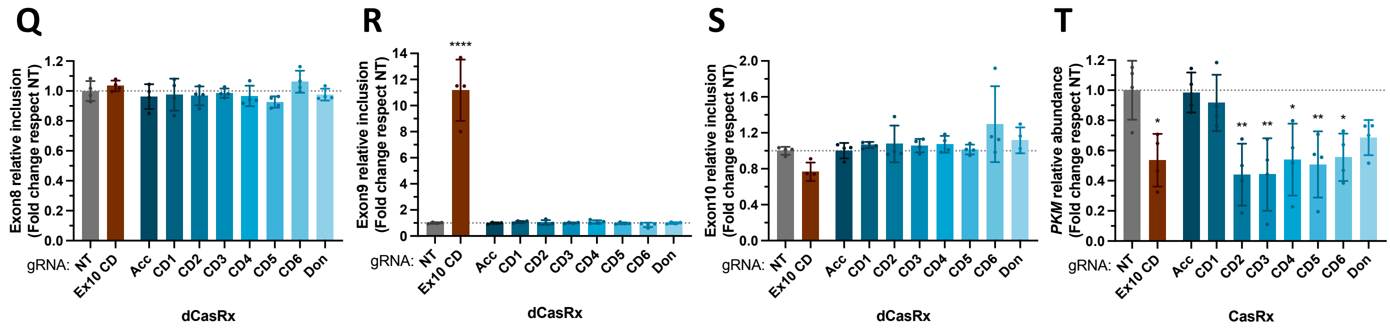


**Supplementary Figure 3. dCasRx position-dependent effect goes beyond the gRNA's targeting efficiency.** (**A**) Genomic location of gRNAs tiled across *PLOD2* exon 14. (**B**) *PLOD2* exon 14 relative inclusion levels in HEK293T cells after transfection of dCasRx and the indicated gRNAs. (**C,D**) *PLOD2* relative RNA abundance after transfection of dCasRx (C) or CasRx (D). (**E**) Genomic location of gRNAs tiled across *ARHGEF11* exon 38. (**F**) *ARHGEF11* exon 38 relative inclusion levels after transfection of dCasRx and the indicated gRNAs. (**G,H**) *ARHGEF11* relative RNA abundance after transfection of dCasRx (G) or CasRx (H). (**I**) Pearson correlation between CasRx RNA cleavage efficiency and dCasRx splice-editing efficiency for each of the gRNAs tiled across *CTNND1*, *SCRIB*, *MBNL1*, *PLOD2*, *ARHGEF11*, *CD46* and *PKM* alternatively spliced exons. (**J**) *CD46* relative RNA abundance after transfection of CasRx and the indicated gRNAs. (**K,P**) Genomic location of gRNAs tiled across *CD46* constitutively spliced exons 4 and 10 (K) or *PKM* exon 8 (P). Acceptor (Acc) and donor (Don) sites were also targeted. (**L-N, Q-S**) Relative exon inclusion levels after transfection of dCasRx and the indicated gRNAs. (**O,T**) Relative RNA abundance after transfection of CasRx. RT-qPCR levels were normalized by total expression of the corresponding gene for splicing analysis; while RNA abundance was normalized by *TBP* housekeeping gene expression. Data were represented as mean ± SD of the fold change relative to non-targeting gRNA (NT) in at least 4 biological replicates. *P <0.05, **P <0.01, ***P <0.001 , ****P <0.0001 in unpaired one-way ANOVA respect NT.

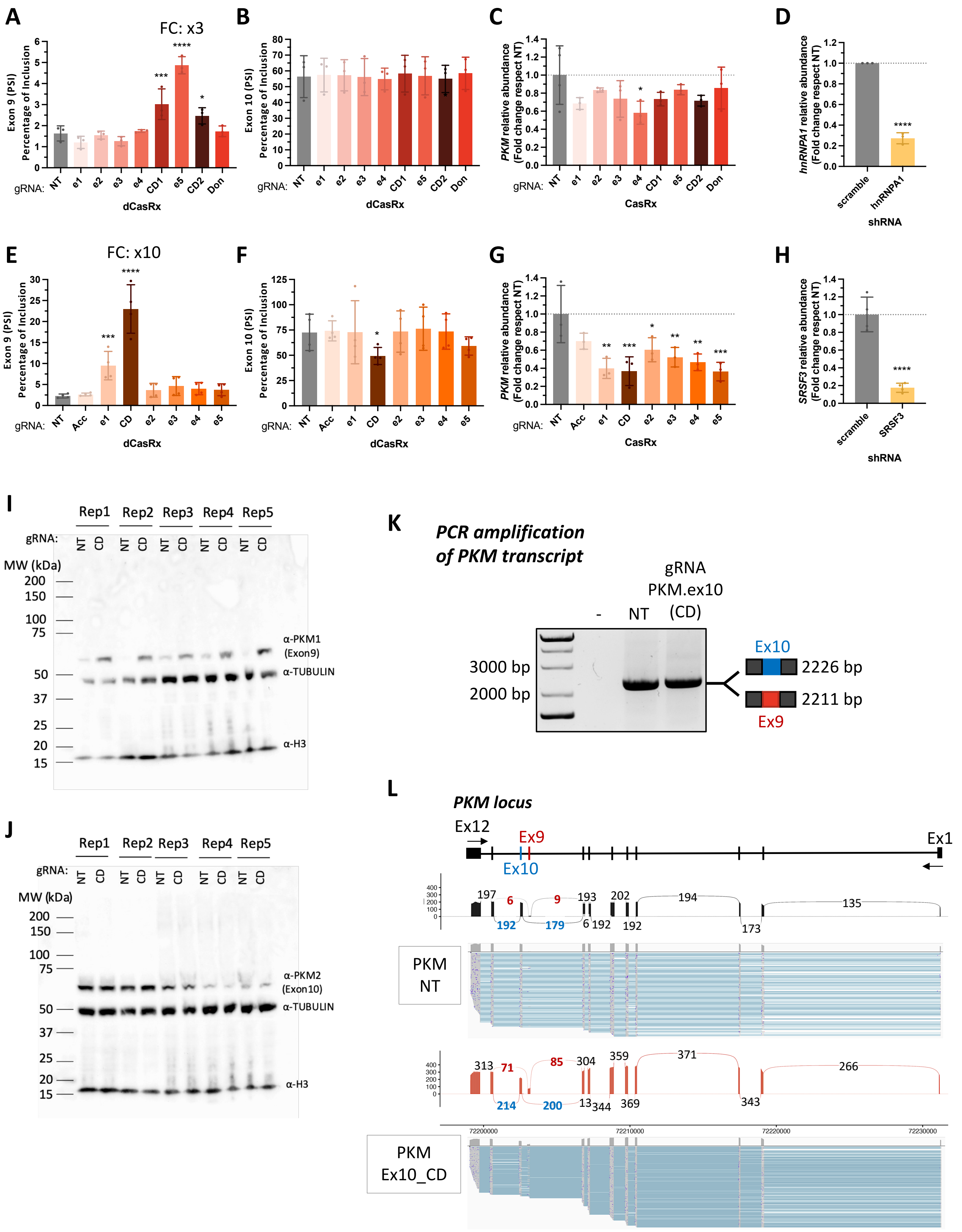
A
FC: x3
Exon 9 (PSI)
Percentage of Inclusion
gRNA: NT e1 e2 e3 e4 CD1 e5 CD2 Don
dCasRx
B
Exon 10 (PSI)
Percentage of Inclusion
dCasRx
C
PKM relative abundance
(Fold change respect NT)
CasRx
D
hnRNPA1 relative abundance
(Fold change respect NT)
scramble hnRNPA1
shRNA
E
FC: x10
gRNA: NT Acc e1 CD e2 e3 e4 e5
F
G
H
SRSF3 relative abundance
(Fold change respect NT)
scramble SRSF3
I
Rep1 Rep2 Rep3 Rep4 Rep5
gRNA: NT CD NT CD NT CD NT CD NT CD
MW (kDa)
α-PKM1
(Exon9)
α-TUBULIN
α-H3
J
α-PKM2
(Exon10)
K
PCR amplification
of PKM transcript
gRNA
PKM.ex10
(CD)
- NT
3000 bp
2000 bp
Ex10
2226 bp
2211 bp
Ex9
L
PKM locus
Ex12
Ex9
Ex10
Ex1
PKM
NT
PKM
Ex10_CD

**Supplementary Figure 4. dCasRx induces accurate splice-switching of *PKM* mutually exclusive isoforms.** (**A-B, E-F**) *PKM* exon 9 and exon 10 percentage of inclusion in HEK293T cells after transfection with dCasRx and the indicated gRNAs targeting *PKM* exon 9 (A,B) or exon 10 (E,F). (**C,G**) *PKM* relative RNA abundance after transfection with CasRx and the indicated gRNAs. (**D,H**) hnRNPA1 and SRSF3 knock-down efficiency. (**I-J**) Uncropped Western Blots detecting PKM1 (**I**) or PKM2 (**J**) isoforms after transfection of dCasRx and the strongest splice-editing gRNA. TUBULIN and H3 were used as loading controls. (**K**) Representative agarose gel showing *PKM* whole transcript after transfection of dCasRx and the strongest splice-editing gRNA. NT is used as a control. Since exon 9 and 10 share the same length, we cannot appreciate a size switch in the gel. (**L**) Whole amplicon sequencing of the RT-qPCR shown in the agarose gel to visualize the changes in splicing across *PKM*. Exons 9 and 10 RT-qPCR levels were normalized by *PKM* total expression levels for splicing analysis; while *PKM* RNA abundance was normalized by *TBP* housekeeping gene expression. Data are represented as mean ± SD of the fold change relative to non-targeting gRNA (NT) in at least 3 biological replicates. *P <0.05, **P <0.01, ***P <0.001, **** P <0.0001 in paired (**A-B, E-F**) or unpaired (**C,G**) one-way ANOVA respect NT and unpaired T-test (**D,H**) respect scramble shRNA.

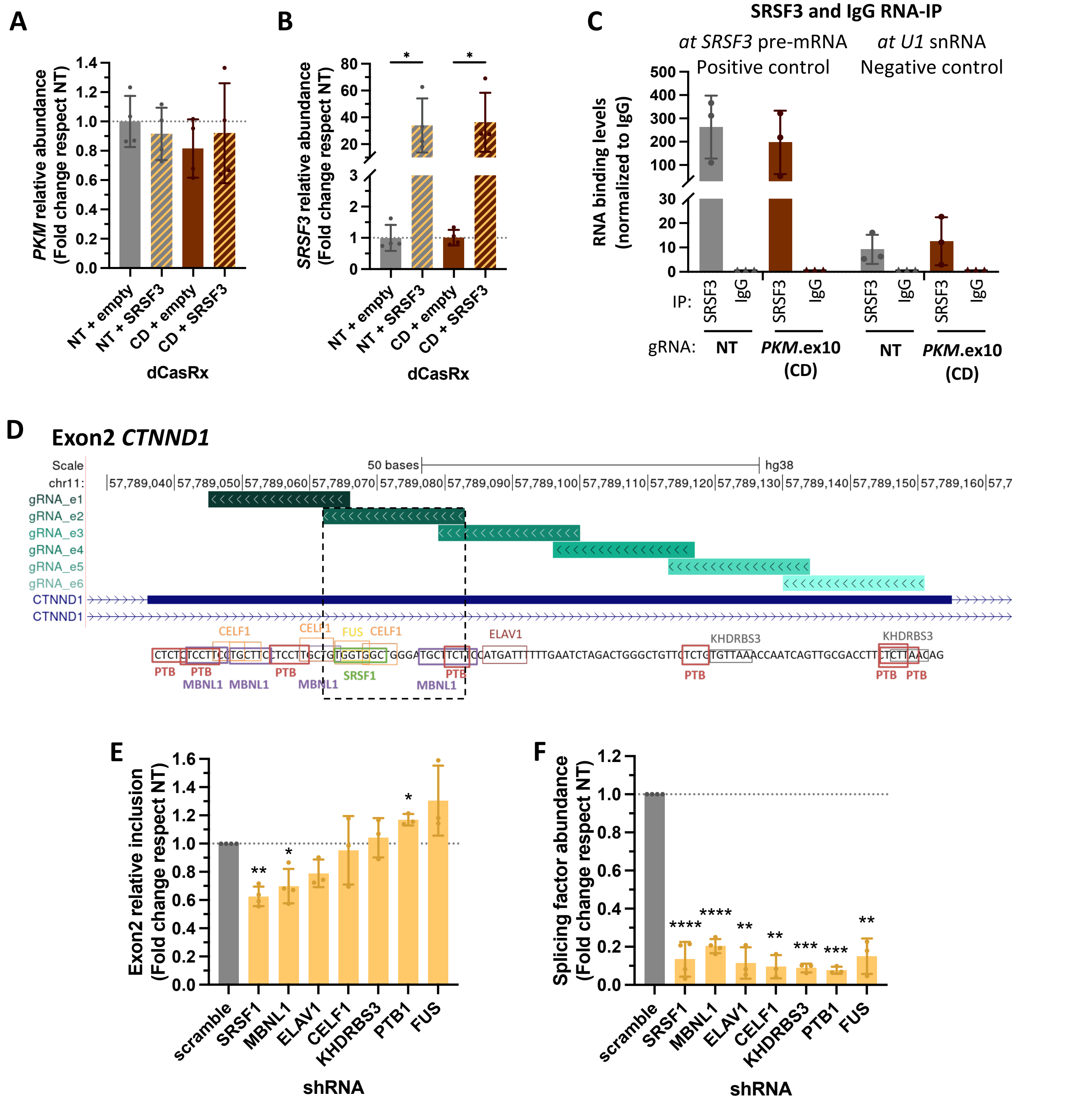


**Supplementary Figure 5. Validation of SRSF3 and SRSF1 mechanistic effects on *PKM* and *CTNND1* splicing.** (**A-B**) *PKM* and *SRSF3* relative RNA levels. (**C**) SRSF3 RNA-IP efficiency by assessing SRSF3 or IgG recruitment levels to control *SRSF3* or *U1* snRNA pre-mRNAs after transfection of HEK293T cells with dCasRx and *PKM's* splice-editing gRNA_CD. % input levels were normalized by IgG recruitment levels in 3 biological replicates. (**D**) Genomic location of *CTNND1* exon 2 targeting gRNAs and the RNA motifs predicted in at least 2 of the 4 RNA databases analysed. The RNA motifs targeted by the strongest splice-editing gRNA are highlighted with a dotted box. (**E**) *CTNND1* exon 2 relative inclusion levels in HEK293T cells after shRNA-mediated knock-down of the indicated splicing factors. (**F**) shRNA knock-down efficiency. Exon 2 RT-qPCR levels were normalized by total *CTNND1* expression levels for splicing analysis; while RNA abundance was normalized to *TBP* housekeeping gene expression. Data are represented as mean ± SD of the fold change relative to NT or scramble shRNA in at least 3 biological replicates. *P <0.05, **P <0.01, ***P <0.001, **** <0.0001 in unpaired one-way ANOVA respect empty (**A-B**) or scramble (**E-F**) .
